# Continuous *ex utero* development of mouse zygotes to early organogenesis

**DOI:** 10.64898/2026.11.20.767777

**Authors:** Hui Yao, Nianqin Sun, Yong Fan, Tianxia Xiao, Kejun Wang, Honglian Shao, Youyue Zhang, Daiji Okamura, Yang Liu, Liling Fan, Junyi Huang, Baiquan Ci, Yi Ding, Tianjin Xia, Zhenyu Yang, Weizhi Ji, Chun So, Bing Bai, Yang Yu, Jun Wu, Tao Tan

## Abstract

In mammals, embryogenesis normally unfolds within the uterus, limiting direct access to the developmental processes that transform a zygote into an embryo with forming organs. Although mouse embryos can be cultured *ex utero* from early post-implantation stages to mid-gestation, continuous culture from the zygote into organogenesis has not been achieved. Through comparative single-cell transcriptomic and metabolomic analyses, here we discover aberrant BMP signaling and suboptimal metabolic support as barriers to continuous *ex utero* development. BMP inhibition, together with stage-specific optimization of the culture environment, enables mouse zygotes to progress through peri-implantation development, gastrulation, and into early organogenesis *ex utero*. Cultured embryos form major embryonic and extraembryonic lineages, establish body axes, and generate multiple organ primordia, while retaining developmental delays and molecular differences from *in vivo* embryos. By connecting developmental stages that have previously been cultured separately, this study provides an accessible platform for dissecting how early mammalian development progresses into organ formation and a reference for evaluating stem-cell-based embryo models.

## Introduction

Mammalian embryogenesis transforms a single-cell zygote into an organized embryo through coordinated changes in cell identity, tissue architecture, signaling and metabolism. Much of this process occurs within the reproductive tract and uterus, making it difficult to observe and manipulate development continuously, particularly across implantation.

*Ex utero* culture has opened important windows into otherwise inaccessible stages of mammalian development. Mouse embryos can be cultured from pre-gastrulation stages (embryonic day (E) 5.5) through organogenesis^1^, whereas human^2–5^ and non-human primate^6,7^ embryos can develop in culture to peri-gastrulation or the onset of organogenesis. These systems, however, begin after several key early developmental events have already occurred *in vivo*. Whether mammalian development can be supported continuously *ex utero* from the zygote into organogenesis therefore remains unresolved. Stem-cell-based embryo models (SCBEMs) offer a complementary approach^8–10^, but they do not reproduce the full development continuum. Some lack essential extraembryonic cell lineages, advanced models generally begin from states corresponding to the peri-gastrulation stage, and mouse blastoids do not support fetal development after uterine transfer. A culture system that bridges pre-implantation development, transition through implantation, gastrulation, and organogenesis would therefore provide direct access to developmental events that are currently studied in separate experimental systems and establish an *ex utero* embryo reference for SCBEMs.

Here we identify elevated BMP signaling and inadequate metabolic support as barriers to continuous *ex utero* development. By combining BMP inhibition and metabolic supplementation with sequential static and roller culture, we support mouse development from the zygote for up to 13 days in culture, reaching features characteristic of early organogenesis and approximately E9.0-E9.5 development.

## Results

### BMP signaling is dysregulated in cultured peri-gastrulation mouse embryos

To identify barriers to continuous development across implantation, we cultured 2-cell mouse embryos (E1.5) in potassium simplex optimized medium (KSOM). Although 93.98% (n = 187) developed to blastocysts within two days, only 6.4% hatched by day 3 (n = 187), approximately when implantation begins *in vivo* (E4.0)^11^ (**Extended Data Fig. 1a-c**). After transfer to Ibidi μ-Slide 8-well plates, only 8.53% formed egg cylinders by day 6 (n = 94; ∼E5.5) (**Extended Data Fig. 1g, h**).

We therefore optimized the pre-implantation culture conditions (**Fig. 1a**, step 1 and **Extended Data Fig. 1a**). From day 2, supplementing KSOM with non-essential amino acids, and sodium pyruvate (referred to as KSOM+) increased hatching to 45.82% by day 3 (n = 218) and increased blastocyst diameter (**Extended Data Fig. 1a-1e**). KSOM+ cultured blastocysts also developed a thickened mural trophectoderm (TE) resembling that of E4.75 embryos *in vivo*^12^ (**Extended Data Fig. 1f)**. Egg cylinder formation increased to 54.38% (n = 113) (**Extended Data Fig. 1g, 1h**). To synchronize subsequent culture, we removed the zona pellucida from unhatched embryos on day 3 and cultured them together with hatched blastocysts in 8-well plates (**Fig. 1a**). Under these conditions, 29.78% of embryos (n = 188) developed to day 8, with morphology and growth comparable to those of *in vivo* E7.25 embryos (**Extended Data Fig. 2a-c**). ZO1 and SOX2 immunofluorescence (IF) staining showed proamniotic cavity formation by day 5 (∼E5.25)^13^. By day 6 (∼E5.5), CER1, LEFTY1 and TOX3 were localized anteriorly, consistent with anterior-posterior axis specification^13,14^. By day 8 (∼E7.25), T expression extended along the posterior primitive streak towards the distal region, where T^+^FOXA2^+^ axial mesoderm cells were detected^15^ (**Extended Data Fig. 2d-f)**. Development then slowed, and embryonic structures deteriorated after transfer to roller culture, collapsing by day 10 (**Extended Data Fig. 2a, b**).

**Fig. 1.**
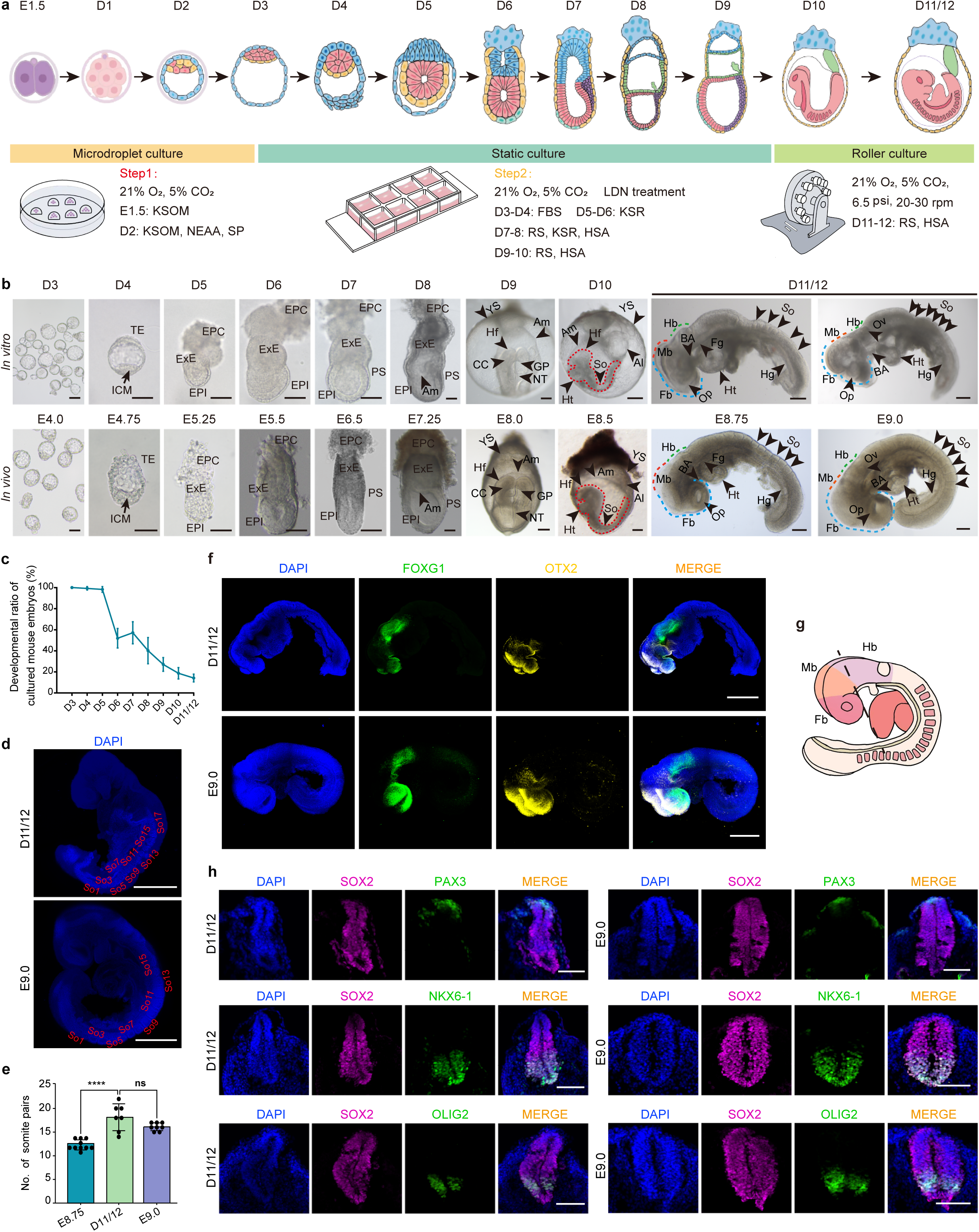
Establishment of an *ex utero* culture system for mouse embryo development from the 2-cell stage to early organogenesis. **a,** Schematic of the *ex utero* culture system for mouse 2-cell embryos. Dx denotes E1.5+Dx. b, Representative brightfield images of cultured embryos (D3-D11/12) and their morphologically equivalent *in viv*o counterparts (E4.0-E9.0). Scale bars, 100 μm for E4.0-E8.5 and D3-D10; 200 μm for E8.75-E9.0 and D11/12. c, Developmental ratio of cultured mouse embryos (n = 244 embryos from 8 independent experimental batches). Data are shown as mean ± S.D. (horizontal lines). d, Representative whole-mount DAPI fluorescence images of cultured D11/12 and *in vivo* E9.0 embryos (n = 7 for D11/12; n = 8 for E9.0). Somites are numbered and their positions are indicated. Scale bar, 500 μm. e, Quantification of somite numbers in E8.75, E9.0, and cultured D11/12 embryos. n = 10, 8, and 7, respectively. Data are shown as mean ± S.D. ns, not significant; ***, *P* < 0.0001 (two-tailed unpaired t-test). **f,** Representative whole-mount immunofluorescence (IF) images of E9.0 and cultured D11/12 embryos stained with DAPI (blue), FOXG1 (green), and OTX2 (yellow), highlighting anterior brain patterning. n = 3 per stage. Scale bar, 500 μm. **g,** Schematic diagram showing the plane of transverse sectioning in cultured D11/12 and *in vivo* E9.0 embryos. **h,** Representative transverse sections of neural tubes from cultured D11/12 and E9.0 embryos, immunostained for pan-neural (SOX2), dorsal (PAX3), and ventral (OLIG2 and NKX6-1) markers. n = 4 for D11/12; n = 3 for E9.0. Scale bar, 100 μm. ICM, inner cell mass; TE, trophectoderm; EPI, epiblast; ExE, extraembryonic ectoderm; EPC, ectoplacental cone; Am, amnion; PS, primitive streak; YS, yolk sac; Hf, head fold; CC, cardiac crescent; GP, gut pocket; NT, neural tube; Al, allantois; Ht, heart; Fb, forebrain; Mb, midbrain; Hb, hindbrain; BA, branchial arches; Fg, foregut; Hg, hindgut; Op, optic pit; Ov, otic vesicle; So, somites.

To investigate this developmental arrest, we performed single-cell RNA sequencing (scRNA-seq) of cultured embryos from days 5-8 (**Supplementary Table 1**) and compared them with published *in vivo* datasets spanning E5.0-E7.25^16,17^. Most major lineages were detected in both *in vivo* and cultured embryos. In contrast, ectoplacental cone cells were identified only in the cultured dataset, as they were not annotated in the *in vivo* reference; epiblast (EPI)-derived non-neural ectoderm was detected only *in vivo* and at very low numbers, suggesting that this population is rare or not maintained in culture (**Extended Data Fig. 3a, b**, and **Supplementary Table 2)**. Gene Ontology enrichment analysis of differentially expressed genes (DEGs) between cultured and *in vivo* embryos showed that BMP, WNT, and TGF-β signaling pathways were commonly enriched among the upregulated DEGs in cultured embryos across multiple cell types (**Extended Data Fig. 3c** and **Supplementary Table 3**). Gene set enrichment analysis further highlighted BMP signaling as a prominent difference between *in vivo* and cultured embryos (**Extended Data Fig. 3d**). Expression levels of BMP receptors (*Bmpr1a*, *Bmpr1b*, *Bmpr2,* and *Acvr1*) and targets (*Id1* and *Id2*) were increased mainly in the epiblast and its derivatives, including the primitive streak (PS) and mesoderm, in cultured embryos (**Extended Data Fig. 3e**). A similar pattern was observed in EPI-like cells from stem cell-based embryo models^18,19^ (**Extended Data Fig. 3f**). Thus, culture from the 2-cell stage is associated with elevated BMP signaling in the EPI and its derivatives, identifying this pathway as a candidate contributor to the observed developmental arrest.

We next tested whether BMP inhibition could extend development. Embryos were treated with LDN193189 (LDN), an inhibitor of BMP type I receptor kinases^20^ (**Fig. 1a**, step 2). LDN reduced *Id1* and *Id2* expression in cultured embryos, including in epiblast and PS cells (**Extended Data Fig. 4a, b**). LDN treatment extended development to days 9 and 10. Day-9 embryos formed a distinct head fold, whereas day-10 embryos displayed somites, forebrain, midbrain and hindbrain regions, a heart tube, foregut and tail bud, broadly resembling E8.0 and E8.5 embryos, respectively (**Extended Data Fig. 4c**). IF staining and three-dimensional (3D) reconstruction of day-10 embryos confirmed HOXB4^+^ somites, SOX2^+^ neuroepithelium, MYH2^+^ myocardium, FOXG1^+^ telencephalon, OTX2^+^ forebrain and midbrain, ISL1^+^ foregut, and CDX2^+^ hindgut (**Extended Data Fig. 4d-h**). The heart tube had begun to loop but remained less developed than in stage-matched *in vivo* embryos (**Extended Data Fig. 4e**). ISL1 was also detected in the heart field, whereas CDX2 also marked the tail bud and posterior neural tube (**Extended Data Fig. 4g, h**)^21^. Thus, BMP inhibition reduces the elevated BMP pathway activity observed in culture and extends *ex utero* development into early organogenesis.

### Continuous development from the 2-cell stage to early organogenesis

We next combined the optimized culture conditions with BMP inhibition. Under these conditions, 2-cell embryos developed *in vitro* for up to 12 days and progressed into early organogenesis (**Fig. 1a-c**). Because embryos at days 11 and 12 reached similar stages, we pooled these time points; 14.17% of starting embryos (n = 244) reached this endpoint. Development was slower and increasingly asynchronous than *in vivo*, particularly after day 9. By days 11/12, 11.76% of embryos had completed axial turning and 47.06% had undergone >50% turning according to established criteria^22^ (**Fig. 1a** and **Extended Data Fig. 5a**).

As axial turning is characteristic of development around E9.0^23^, we compared day-11/12 cultured embryos with E9.0 embryos *in vivo*. Cultured embryos showed ongoing neural tube (NT) closure, visible optic vesicles and the auditory pit, and 14-22 somite pairs, overlapping with the range at E9.0 (**Fig. 1b**, **d**, **e**). SOX9^+^ and TWIST^+^ domains were present within developing somites^24,25^ (**Extended Data Fig. 5b**). The nervous system also showed regional patterning: FOXG1 marked the telencephalon and other cranial domains^26^, whereas FOXA2, NKX2-2, OLIG2, and NKX6-1 marked ventral domains, and PAX3 marked the dorsal domain of the SOX2-positive NT (**Fig. 1f-h** and **Extended Data Fig. 5c**), consistent with dorsal-ventral patterning at E9.0-E9.5^27–29^. Cultured embryos also formed a C-shaped looping heart, with TBX5^+^ and ISL1^+^ domains corresponding to first- and second-heart-field derivatives^30^ (**Extended Data Fig. 5d, f**). These morphological and molecular features indicate that embryos cultured continuously from the 2-cell stage can progress through gastrulation and acquire features of early organogenesis approaching an E9.0-like stage.

### *Ex utero* embryos reproduce major cellular and regulatory features of *in vivo* development

To define cell states and gene regulatory programs during *ex utero* development, we performed scRNA-seq and single-cell assay for transposase-accessible chromatin using sequencing (scATAC-seq) on cultured embryos. To account for developmental asynchrony after day 9, cultured embryos were selected by morphology: embryos corresponding approximately to E8.0, E8.5, and E9.0 were collected on days 9-10, 10-11, and 11-12, respectively (**Supplementary Table 1**). Because suitable published chromatin-accessibility data were not available for all comparison stages, we also generated scATAC-seq profiles from stage-matched *in vivo* embryos. scRNA-seq analysis revealed progressive diversification of cell types from days 5 to 12 (**Fig. 2a**, **Supplementary Table 2, and Extended Data Fig. 6a, d**). Before day 8, extraembryonic lineages comprised a substantial fraction of cells, including TE, primitive endoderm (PrE), and their derivatives. PS cells emerged by day 7, followed by the expansion and diversification of mesodermal and endodermal lineages from day 8. By days 9/10, progenitor populations associated with early organogenesis appeared, followed by further diversification of neural and other organ-associated cell types at days 11/12.

**Fig. 2.**
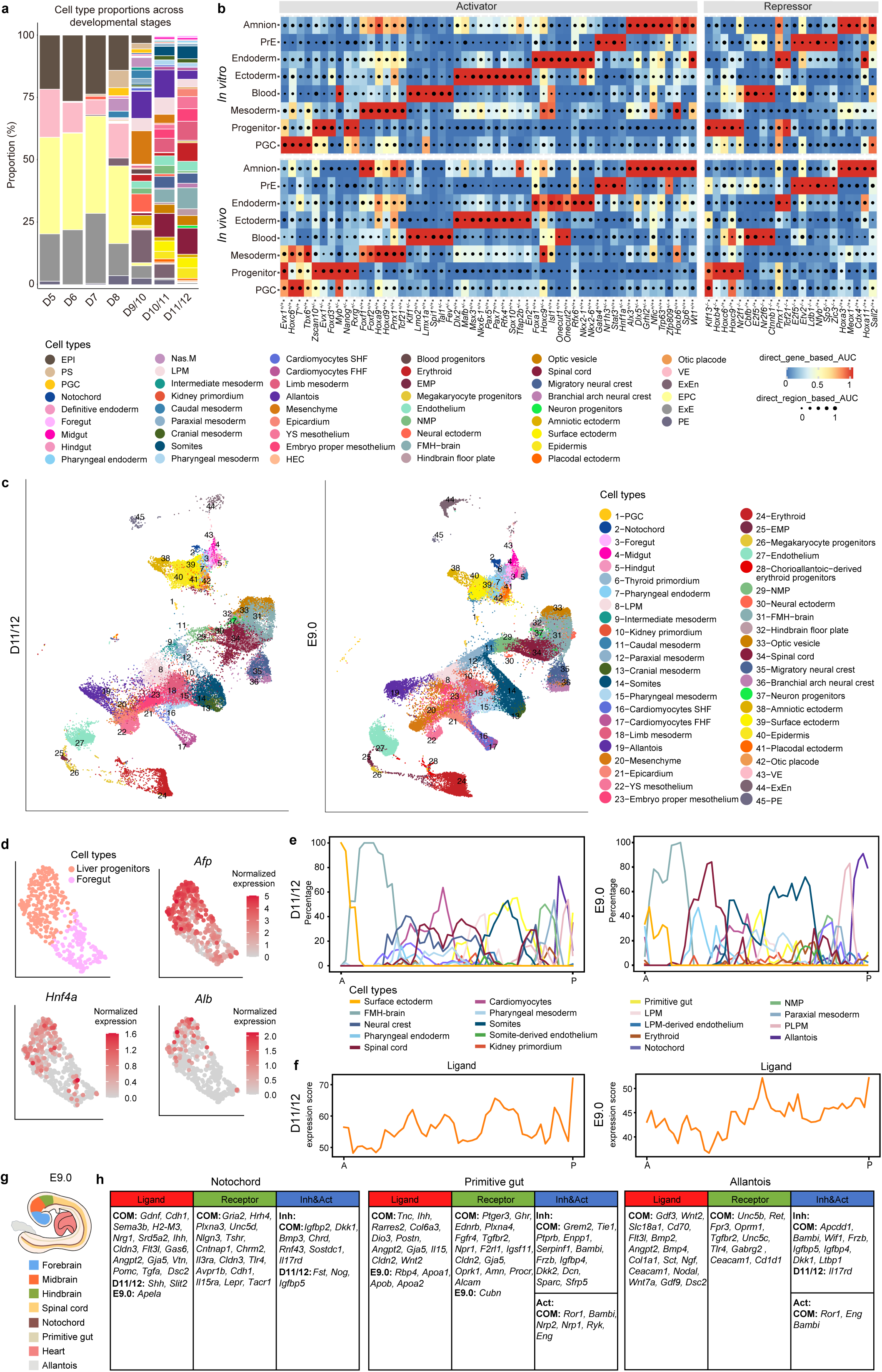
Comparative analysis of single-cell multi-omics and spatial transcriptomic profiles of cultured and *in vivo* embryos. **a,** Stacked bar plot showing cell type proportions across *in vitro* developmental stages (D5-D11/12; Dx denotes E1.5+Dx). Each color represents a distinct cell type. **b,** SCENIC+ analysis showing major lineage-specific regulons (eRegulons) across cultured and *in vivo* embryos. Color indicates direct gene-based AUC activity, while dot size represents direct region-based AUC activity. **c,** UMAP visualization of integrated scRNA-seq datasets from cultured D11/12 (left) and *in vivo* E9.0 (right) embryos. Each dot represents one cell, and colors encode the cell type. **d,** UMAP plots showing liver progenitor cells within the foregut cluster from the cultured D11/12 scRNA-seq dataset. The upper-left panel shows the UMAP projection with cell types color-coded. The upper-right, lower-left, and lower-right panels show feature plots of the expression of liver marker genes *Afp*, *Hnf4a*, and *Alb*, respectively, with color intensity indicating gene expression levels. **e,** Cell type proportions along the embryonic anterior-posterior (A-P) axis of cultured D11/12 (left) and *in vivo* E9.0 (right) embryos. Colors denote distinct cell types. **f,** Expression score of CellChat-derived ligands across cells within sliding windows along the A-P axis of cultured D11/12 and *in vivo* E9.0 embryos, based on Stereo-seq spatial transcriptomic datasets. **g,** Schematic illustration of major signaling centers in E9.0 mouse embryos (lateral view). Colors denote distinct cell types/tissues. **h,** Comparison of major signaling centers between cultured D11/12 embryos and *in vivo* E9.0 embryos. “COM” denotes signaling genes with comparable expression levels between the two datasets (adjusted *P* value ≥ 0.05). “D11/12” and “E9.0” denote genes with significantly higher expression in cultured D11/12 or *in vivo* E9.0 embryos, respectively (adjusted *P* value < 0.05 and log2FC > 1). EPI, epiblast; PS, primitive streak; PGC, primordial germ cells; Nas.M, nascent mesoderm; LPM, lateral plate mesoderm; Cardiomyocytes SHF, second heart field-derived cardiomyocytes; Cardiomyocytes FHF, first heart field-derived cardiomyocytes; YS mesothelium, yolk sac mesothelium; HEC, haemato-endothelial cells; EMP, erythroid-myeloid progenitors; NMP, neuro-mesodermal progenitors; FMH-brain, forebrain/midbrain/hindbrain; VE, visceral endoderm; ExEn, extraembryonic endoderm; EPC, ectoplacental cone; ExE, extraembryonic ectoderm; PE, parietal endoderm; PLPM, posterior lateral plate mesoderm; PrE, primitive endoderm; Inh, inhibitor; Act, activator.

To compare gene regulatory networks between cultured and *in vivo* embryos, we annotated scATAC-seq cell types (**Extended Data Fig. 6b, c, e**) and then integrated the scRNA-seq and scATAC-seq data to perform SCENIC+ analysis^31^. Major lineage-associated regulons showed broadly similar activity in the two conditions. For example, endoderm-associated regulons including *Foxa1*, *Onecut1*, *Nkx2-1,* and *Nkx2-6* showed similar lineage-specific activity. Differences were also evident, including lower activity of the mesoderm-associated *Tbx6* and *T* regulons in cultured embryos than in E9.0 embryos (**Fig. 2b**).

We next determined the developmental stage of *ex utero* day-11/12 embryos. Integrated transcriptomic analysis showed that their major cell populations aligned more closely with *in vivo* E9.0 embryos than with E8.5 or E8.75 embryos **(Extended Data Fig. 6f**). We identified 37 embryonic and 6 extraembryonic cell types based on established markers^17^ (**Supplementary Table 2**), most with corresponding populations in E9.0 embryos. Interestingly, intermediate mesoderm was only detected in day-11/12 embryos, whereas thyroid primordium and chorioallantoic-derived erythroid progenitors were only found in E9.0 embryos *in vivo* (**Fig. 2c**), indicating incomplete or delayed specification of some lineages. Other foregut-derived progenitors were present: a subset of pharyngeal endoderm cells expressed *Nkx2-1* together with *Irx2*, *Gata4/6* and *Cpm*, consistent with an early lung progenitor identity^32^ (**Extended Data Fig. 6g**) and a distinct foregut population expressed *Hnf4a*, *Afp* and *Alb*, consistent with hepatic specification^33^ (**Fig. 2d**). Thus, day-11/12 embryos reproduce much of the cellular diversity of E9.0 embryos while differing in the representation or maturation of specific lineages.

We next examined whether spatial signaling organization was maintained during *ex utero* organogenesis. Stereo-seq analysis of mid-sagittal sections from day-11/12 cultured and E9.0 *in vivo* embryos identified 19 major cell types with broadly comparable spatial distributions (**Extended Data Fig. 7a-e** and **Supplementary Table 1, 2**). Regions along the anterior-posterior axis with high cell density and enriched ligand expression were defined as putative signaling centers^34^. The primitive gut, notochord, and allantois met these criteria in both cultured and *in vivo* embryos (**Fig. 2e, f,** and **Extended Data Fig. 7f**). Comparison of CellChat-defined ligands, receptors, activators, and inhibitors^35^ showed that most detected signaling components were shared between the two conditions, although specific differences remained. Cultured embryos showed higher *Shh* expression in the notochord and *Il17rd* in the allantois, whereas *Cubn* was higher in the *in vivo* primitive gut (**Fig. 2g, h**).

As an example of predicted communication between signaling centers and adjacent tissues, we examined the primitive gut. Many inferred pathways were shared between day-11/12 and E9.0 embryos (**Extended Data Fig. 7g**). *Lama1* and *Lamb5* were expressed in the primitive gut, whereas their predicted receptors *Dag1* and *Itga6* were expressed in cardiomyocytes in both conditions, suggesting a conserved role for gut-derived LAMININ in extracellular matrix remodeling and cardiac morphogenesis^36^ (**Extended Data Fig. 7h**). Other pathways, including WNT, UNC5 and HH signaling, differed between cultured and *in vivo* embryos (**Extended Data Fig. 7g**). Together, these analyses show that *ex utero* embryos reproduce major cellular, gene-regulatory and spatial signaling features of E9.0 development while retaining differences in specific lineages and signaling programs.

### Metabolomic remodeling during *ex utero* development

Metabolism changes substantially during embryogenesis and is closely linked to developmental progression^37^. We therefore asked whether embryos developing *ex utero* undergo the metabolic transitions observed *in vivo*. We profiled metabolites in *in vivo* embryos from E5.5 to E9.0 and cultured embryos from days 6 to 12. For stage-matched analysis, cultured embryos were grouped according to morphology, with E8.0-, E8.5-, and E9.0-equivalent embryos collected on days 9-10, 10-11, and 11-12, respectively. After data processing, 261 annotated metabolites were retained for analysis (**Extended Data Fig. 8**).

Correlation analysis showed that cultured embryos shared stage-dependent metabolic features with their *in vivo* counterparts (**Fig. 3a**). Day-11/12 embryos, for example, were more similar to E9.0 than to E8.75 embryos. Both cultured and *in vivo* embryos showed a marked metabolic transition between late gastrulation and early organogenesis, around E7.25-E7.75 *in vivo* and days 8-9 in culture (**Fig. 3a**). Principal component analysis also revealed important differences. PC1 captured the developmental progression of *in vivo* embryos from E5.5 to E9.0, with a prominent transition around E7.75-E8.0. Cultured embryos followed a similar trajectory until around day 9 but showed less progression thereafter. PC2 separated cultured from *in vivo* embryos, indicating a metabolic effect of the *ex utero* environment, whereas PC3 revealed additional differences in their developmental trajectories (**Fig. 3b**). Consistent with the correlation analysis, Euclidean distances between consecutive stages identified a prominent metabolic transition between E7.25 and E7.75 *in vivo* and between days 8 and 9 in culture (**Fig. 3c)**. Thus, cultured embryos undergo a metabolic shift at a similar developmental stage to *in vivo* embryos, but their later metabolic trajectories diverge.

**Fig. 3.**
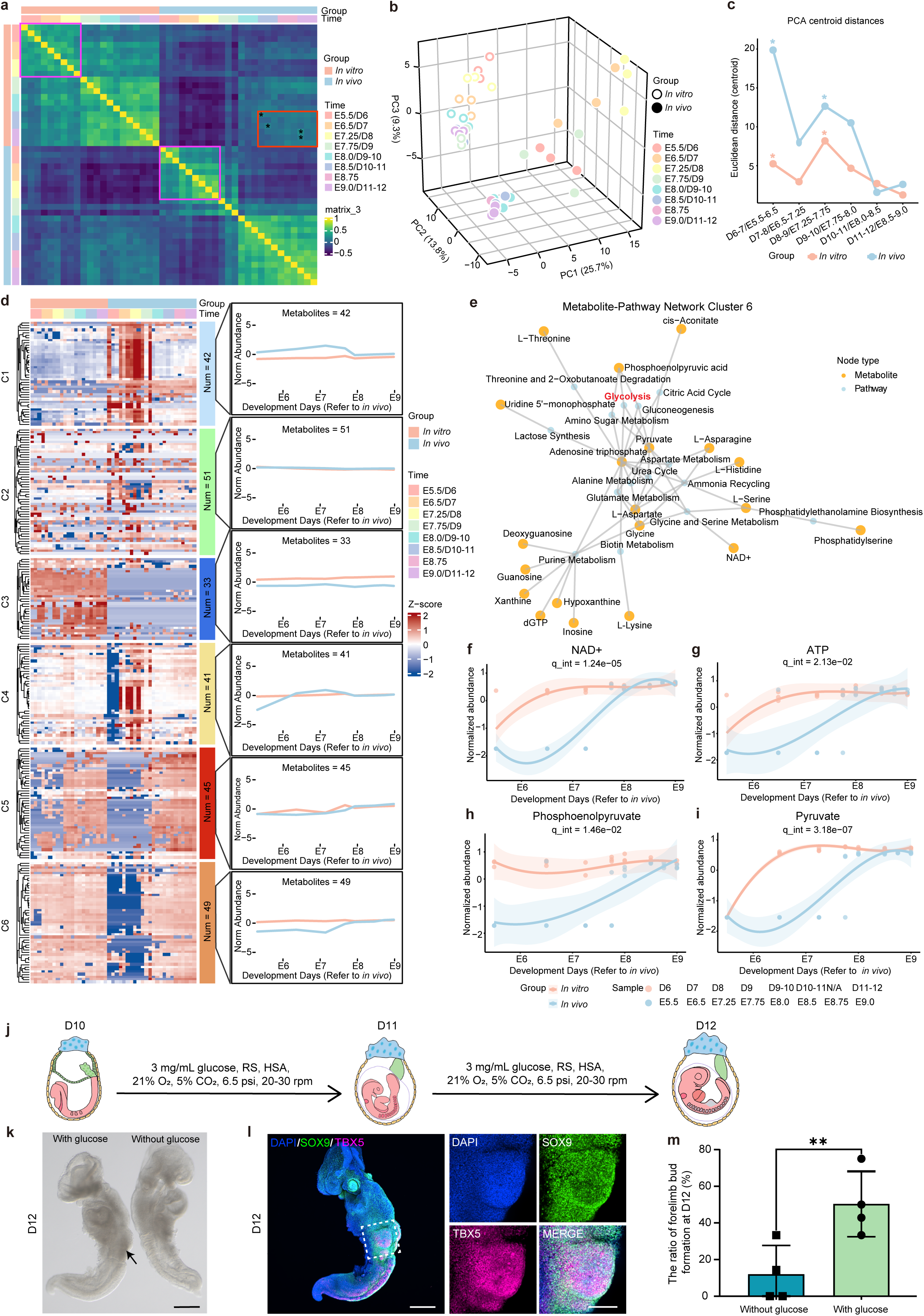
Metabolic dynamics during *in vitro* embryogenesis. **a,** Heatmap of Spearman’s correlation coefficients between *in vivo* (E5.5-E9.0) and cultured (D6-D12, Dx denotes E1.5+Dx) embryos based on 261 annotated metabolites. Magenta squares indicate stages before E7.25 (*in vivo*) or D8 (*in vitro*). The red square marks the correlation pattern in which most D10-D11 samples correlate most strongly with E8.5, whereas most D11-D12 samples correlate most strongly with E9.0 (* denotes the most highly correlated *in vitro*-*in vivo* sample pairs within the red square). **b,** 3D PCA plot showing all *in vivo* and *in vitro* samples. Different colors indicate distinct developmental stages; solid and open circles represent *in vivo* and *in vitro* samples, respectively. **c,** Line plot showing PCA centroid distances between *in vivo* and *in vitro* samples. * indicates the two most pronounced changes in centroid distance, occurring between D6-D7/E5.5-E6.5 and D8-D9/E7.25-E7.75, respectively. **d,** Heatmap showing the normalized abundance (Z-score) of 261 annotated metabolites across *in vivo* and *in vitro* embryos over developmental stages. Metabolites are grouped into six clusters, with the average abundance dynamics shown for each cluster. **e,** Network of significantly enriched pathways (blue solid circles) and their associated metabolites (yellow solid circles) in Cluster 6. **f,** Normalized abundance of NAD^+^ across developmental stages (*in vivo* E5.5-E9.0; *in vitro* D6-D12), fitted with cubic polynomial regression (solid lines), with shaded areas representing the 95% confidence intervals. The interaction term (q_int) between developmental stages and groups is significant (*P* < 0.0001). **g,** Normalized abundance of ATP changing across developmental stages (*in vivo* E5.5-E9.0; *in vitro* D6-D12), fitted with cubic polynomial regression (solid lines), with shaded areas representing the 95% confidence intervals. The interaction term (q_int) between developmental stages and groups is significant (*P* < 0.05). **h,** Normalized abundance of phosphoenolpyruvate across developmental stages (*in vivo* E5.5-E9.0; *in vitro* D6-D12), fitted with cubic polynomial regression (solid lines), with shaded areas representing the 95% confidence intervals. The interaction term (q_int) between developmental stages and groups is significant (*P* < 0.05). **i,** Normalized abundance of pyruvate across developmental stages (*in vivo* E5.5-E9.0; *in vitro* D6-D12), fitted with cubic polynomial regression (solid lines), with shaded areas representing the 95% confidence intervals. The interaction term (q_int) between developmental stages and groups is significant (*P* < 0.0001). **j,** Schematic of glucose supplementation in the 2-cell embryo culture system. **k,** Representative brightfield images of D12 embryos from glucose-supplemented (n = 21) and glucose-free (n = 20) culture conditions. The arrow indicates the forelimb bud. Scale bar, 500 μm. **l,** Representative whole-mount IF images of D11/12 embryos cultured in glucose-supplemented medium, stained with DAPI (blue), TBX5 (magenta), and SOX9 (green), highlighting forelimb bud formation. Scale bars, 500 μm (left) and 100 μm (right). **m,** Bar plot showing the percentage of D12 embryos with forelimb buds following culture in glucose-supplemented or glucose-free medium. Data are mean ± S.D. \*\**P* < 0.01; n = 21 for the glucose-supplemented group and n = 20 for the glucose-free group, from 4 independent batches.

To gain insight into the metabolic differences between *ex utero* and *in vivo* embryos, we grouped the 261 metabolites into six clusters based on their developmental trajectories (**Fig. 3d** and **Supplementary Tables 4, 5**). Cluster 1 contained metabolites that were generally more abundant *in vivo*, particularly before E7.75, and this cluster was enriched for arginine and proline metabolism (**Extended Data Fig. 9a**). This cluster included creatine, fumarate, taurine, and several acylcarnitines (**Extended Data Fig. 9b-g**), indicating differences in nutrient availability and lipid metabolism between the two environments. Clusters 2 and 3 showed relatively limited developmental changes; many metabolites enriched in Cluster 3 in cultured embryos were components or derivatives of the culture environment, such as ethylene glycol derivatives (**Fig. 3d** and **Supplementary Table 4**). Cluster 4 contained metabolites that changed around E7.25-E8.0 *in vivo* but remained comparatively stable in culture and was enriched for several amino acid metabolic pathways (**Fig. 3d** and **Extended Data Fig. 9h**). This cluster included α-ketoglutarate and succinate, together with glutamine, glutamate, and phosphate (**Extended Data Fig. 9j-n** and **Supplementary Table 4**), suggesting a loss of the metabolic remodeling of the TCA cycle under *ex utero* culture conditions. By contrast, Cluster 5 captured metabolic changes that were largely preserved in culture. These metabolites increased with development, with a prominent rise between E7.25 and E7.75, and were enriched for glycolysis, the pentose phosphate pathway, and other central carbon metabolic pathways (**Fig. 3d** and **Extended Data Fig. 10a**). This glycolytic upshift was also reflected at the transcriptomic level, as evidenced by the increased mRNA expression of glycolysis-related enzymes (**Extended Data Fig. 10b**). Several glycolytic and related metabolites, including glucose-6-phosphate, fructose-6-phosphate, dihydroxyacetone phosphate, ribose-5-phosphate, glycerol-3-phosphate, and lactate, showed broadly similar developmental trajectories *in vivo* and *in vitro* (**Extended Data Fig. 10c-h**). Cluster 6, however, revealed a metabolic transition that was not reproduced in culture. These metabolites increased sharply between E7.25 and E7.75 *in vivo* but remained relatively stable in cultured embryos (**Fig. 3d**). The cluster was enriched for glycolysis/gluconeogenesis and the citric acid cycle (**Fig. 3e**). NAD+ also followed different trajectories *in vivo* and *in vitro* (**Fig. 3f**). Notably, ATP, phosphoenolpyruvate, and pyruvate increased around E7.75 *in vivo*, whereas this increase was absent in cultured embryos despite their relatively high basal levels (**Fig. 3g-i**). Together, these data indicate that cultured embryos reproduce some metabolic changes associated with early organogenesis but fail to undergo the full metabolic remodeling observed *in vivo*.

Given the differences in glycolytic metabolites during this transition, we tested whether increasing glucose availability could improve later *ex utero* development. Additional glucose was supplied from day 10 (**Fig. 3j**). Glucose supplementation increased the frequency of embryos with morphologically identifiable forelimb buds, which were confirmed by SOX9 and TBX5 staining (**Fig. 3k-l**). The proportion of embryos forming forelimb buds was significantly higher with glucose supplementation than under the control condition (*P* < 0.01; **Fig. 3m**). Thus, the metabolomic comparison identified a metabolic divergence during early organogenesis and provided a basis for optimizing culture conditions to further improve *ex utero* mouse embryonic development.

### *Ex utero* embryogenesis from the zygote to early organogenesis

We next asked whether the optimized culture system could support continuous development starting directly from zygotes. Because zygotes require approximately one additional day to reach the 2-cell stage, transfer to roller culture were shifted by one day, to day 11. After three days in KSOM followed by one day in KSOM+, 85.42% of zygotes (n = 294) reached the blastocyst stage, compared with 94.09% of embryos cultured from the 2-cell stage (n = 274). Development declined markedly between days 6 and 7, and only 5.62% of the starting embryos reached day 12 (**Extended Data Fig. 11a**).

Despite this reduced efficiency, embryos that continued developing underwent the expected sequence of major developmental events. At peri-implantation stages, OCT4^+^ EPI, CDX2^+^ TE and GATA6^+^ PrE lineages were detected, followed by lumen formation within the EPI (**Extended Data Fig. 11b**). A-P polarity subsequently emerged, with DKK1 localized anteriorly and T posteriorly (**Extended Data Fig. 11c**). During gastrulation, the three germ layers became regionally organized. OTX2^+^ cells were positioned anteriorly next to FOXA2^+^ anterior visceral endoderm, consistent with specification of anterior ectoderm^38^. T^+^/FOXA2^+^/OTX2^+^ cells were also detected in the anterior PS, consistent with molecular features of the organizer region^39–42^ (**Extended Data Fig. 11d**). Thus, embryos cultured from zygotes can undergo key lineage specification and patterning events, although with reduced developmental efficiency.

By day 12, surviving embryos displayed morphological features broadly comparable to E8.75 embryos (**Extended Data Fig. 11e**). They contained 12-14 bilaterally arranged somite pairs (**Extended Data Fig. 11f, g**) and formed an organized primitive gut. ISL1^+^ anterior and CDX2^+^ posterior domains were consistent with the foregut and hindgut development, respectively^43,44^ (**Extended Data Fig. 11h**). Differences from *in vivo* development were also apparent. SOX2^+^OCT4^+^ primordial germ cells (PGCs) were detected more anteriorly within the hindgut of cultured embryos, whereas most remained at more posterior positions in stage-matched *in vivo* embryos (**Extended Data Fig. 11i**), suggesting altered timing or positioning of PGC migration.

Because our metabolomic analysis identified differences in glycolysis between cultured and *in vivo* embryos during the transition to organogenesis, we next tested whether supplementation with human umbilical cord blood serum (HCS), which contains glycolysis-promoting factors, including glucose^45^, PDGF-BB^46,47^, and IGF1^48^, could improve later development. Embryos received LDN treatment, followed by HCS (**Fig. 4a**). Upon neural-fold formation on day 11, embryos were transferred to roller culture (**Fig. 4a**). HCS supplementation increased developmental efficiency after day 7 and extended development to day 13 (**Fig. 4b, c**). Day-13 embryos underwent axial turning and formed multiple organ primordia, including optic and auditory structures, heart and primitive gut (**Fig. 4d** and **Supplementary Video 1**). Two morphologically distinct branchial arches were also present, a feature characteristic of development around E9.5^49^ (**Fig. 4d**). Consistent with this staging, day-13 embryos contained 20-28 somite pairs, overlapping with the range of E9.5 embryos and exceeding that of E9.0 embryos (**Fig. 4e**). IF staining further confirmed organization of several developing tissues. A cup-shaped epithelial structure posterior to the brain expressed SOX2 and SOX9, consistent with the developing auditory vesicle^50^ (**Fig. 4f**). DLX2^+^ cells marked two branchial arches adjacent to the cranial region^51^ (**Fig. 4g**). CD31^+^ endothelial cells^52^ formed a vascular network surrounding TUJ1^+^ neural tissue^53^, consistent with formation of the perineural vascular plexus^54^. Intersomitic vessels were also detected extending dorsally from the dorsal aorta, resembling the vascular organization observed at this stage *in vivo*^55^ (**Fig. 4h** and **Supplementary Video 2**). Together, these results establish culture conditions that support continuous mouse development *ex utero* from the zygote through peri-implantation development, gastrulation and into early organogenesis. HCS supplementation further extends this developmental trajectory and improves the formation of organ primordia, although developmental efficiency remains substantially lower than *in vivo*.

**Fig. 4.**
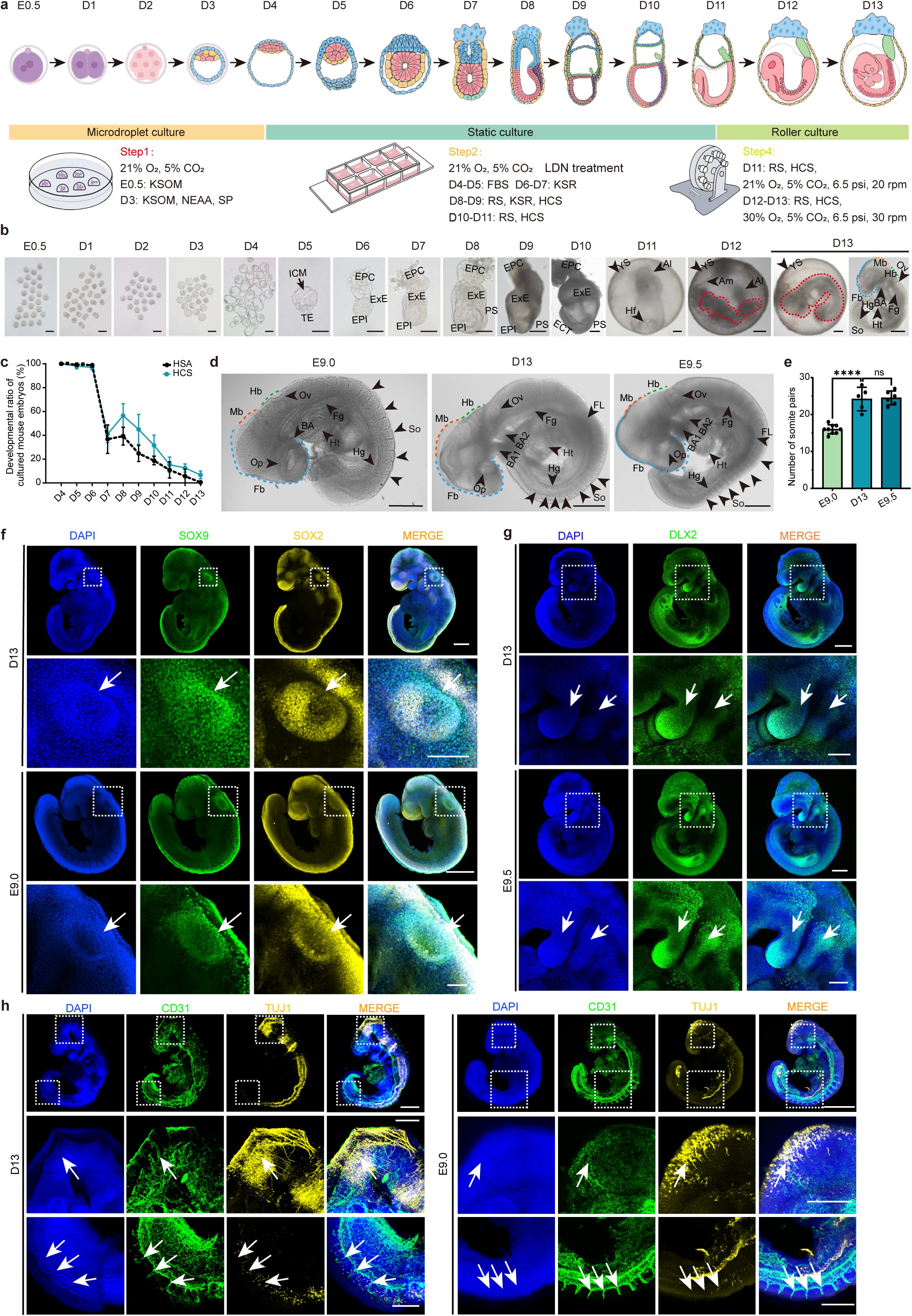
Extending the *ex utero* culture system from the zygote to early organogenesis. **a,** Schematic of the optimized mouse embryo culture system from zygote to early organogenesis (Dx denotes E0.5+Dx). **b,** Representative brightfield images of cultured embryos from E0.5 to D13. Scale bars, 100 μm (E0.5-D8), 200 μm (D9-D11), and 500 μm (D12-D13). **c,** Developmental ratio of cultured embryos from D4 to D13 in media supplemented with human serum albumin (HSA) or human umbilical cord serum (HCS). HSA group, n = 205 embryos from 7 independent experiments; HCS group, n = 100 embryos from 4 independent experiments. Data are presented as mean ± S.D. **d,** Representative bright-field images of cultured D13 embryos alongside *in vivo* E9.0 and E9.5 embryos. Morphologically evident structures (e.g., heart, somites, and brain) are labeled. Scale bar, 500 μm. **e,** Quantification of somite numbers in E9.0, cultured D13, and E9.5 embryos (n = 9, 5, and 6, respectively). Data are presented as mean ± S.D. ns, not significant; ***, *P* < 0.0001 (two-tailed unpaired t-test). **f,** Representative whole-mount IF images of cultured D13 embryos and *in vivo* E9.0 embryos stained with DAPI (blue), SOX9 (green), and SOX2 (yellow). The lower panels show higher-magnification views of the boxed regions in the upper panels, with white arrows indicating the otic vesicle. n = 3 for D13 embryos; n = 3 for E9.0 embryos. Scale bars, 500 μm (upper) or 100 μm (lower). **g,** Representative whole-mount IF images of cultured D13 embryos and *in vivo* E9.5 embryos stained with DAPI (blue) and DLX2 (green). The lower panels show higher-magnification views of the boxed regions in the upper panels, with white arrows indicating the branchial arches. n = 3 for D13 embryos and n = 3 for E9.5 embryos. Scale bars, 500 μm (upper) or 200 μm (lower). **h,** Representative whole-mount IF images of cultured D13 embryos and *in vivo* E9.0 embryos stained with DAPI (blue), CD31 (green), and TUJ1 (yellow). The middle panels show higher-magnification views of the boxed regions in the brain, highlighting the perineural vascular plexus (white arrows) surrounding the neural cells. The lower panels show magnified views of the boxed regions in the somites, with white arrows indicating intersomitic vessels. n = 5 for D13 embryos and n = 4 for E9.0 embryos. Scale bars, 500 μm (upper) or 200 μm (middle and lower). ICM, inner cell mass; TE, trophectoderm; EPI, epiblast; ExE, extraembryonic ectoderm; EPC, ectoplacental cone; ECT, ectoderm; Am, amnion; Al, allantois; YS, yolk sac; Hf, head fold; Fb, forebrain; Mb, midbrain; Hb, hindbrain; BA, branchial arches; Fg, foregut; Hg, hindgut; Ht, heart; Op, optic pit; Ov, otic vesicle; So, somites.

## Discussion

Through stepwise optimization, here we established a culture system that supports continuous *ex utero* development of mouse embryos from the zygote to early organogenesis. This required stage-specific interventions at different timepoints: improved support for pre-implantation development, inhibition of BMP signaling during peri-gastrulation development, and modification of the metabolic environment during early organogenesis. Under these conditions, embryos progressed through major developmental transitions, including embryonic and extraembryonic lineage specification, anterior–posterior patterning, gastrulation, and the formation of early organ primordia, including the brain, heart, and gut. By bridging pre- and post-implantation development, which have traditionally required separate culture regimes^56^, our approach provides continuous experimental access to a period of mammalian development that normally unfolds within the reproductive tract and uterus.

Our findings also identify barriers that emerge when mammalian development is removed from the maternal environment. Comparisons with stage-matched *in vivo* embryos revealed altered BMP signaling during peri-gastrulation development and metabolic differences during the transition to organogenesis. BMP inhibition and subsequent modification of the culture environment extended development, illustrating how systematic molecular comparison with *in vivo* embryos can guide improvements in *ex utero* culture. Nevertheless, important differences remain. Development was slower and less efficient than *in vivo*, became increasingly asynchronous at later stages, and did not progress beyond early organogenesis. The cultured embryos also showed differences in specific cell populations, gene-regulatory programs and metabolic states. Defining which maternal, extraembryonic and metabolic inputs become limiting at later stages will therefore be important for extending development further.

More broadly, continuous development from a zygote into organogenesis represents an initial step towards ectogenesis in a mammalian species whose embryonic development normally depends on the maternal reproductive tract. Full ectogenesis would require support for the entire developmental trajectory to a viable organism, which remains far beyond what is achieved here. Nevertheless, extending *ex utero* development across implantation, gastrulation, and into organogenesis begins to turn a normally inaccessible developmental continuum into an experimentally accessible process. This framework will enable direct study of how early events influence later organ formation, provide an *ex utero* reference for evaluating stem-cell-based embryo models, and offer a path towards progressively extending *ex utero* development in mammals.

## Method

### Ethical statement

The animal experiments were approved by the Experimental Animal Welfare and Ethics Committee of Kunming University of Science and Technology (Approval No. PZWH (DIAN) 2020-0003). Human umbilical cord blood was collected from healthy pregnant women at the prenatal clinics of Peking University Third Hospital and the Second Affiliated Hospital of Kunming Medical University, with informed consent obtained from all participants. The study was approved by the Ethics Committees of both institutions (Approval No. 780-04 and PJ-KY-2026-313).

### Animal

Adult male and female ICR mice, 8–10 weeks of age, were purchased from Beijing Sibefu Biotechnology Co., Ltd. All mice were housed at 21.5–23 °C and 30–50% humidity under a 12-h light/12-h dark cycle.

### Superovulation and embryo retrieval

8- to 10-week-old CD1 female mice were injected intraperitoneally with 10 IU/mouse PMSG (Ningbo Second Hormone Factory) in saline (0.9% NaCl) (Shaanxi Sheng’ao). After 46-48 h, the mice were injected with 10 IU/mouse hCG (Ningbo Second Hormone Factory) and mated with males. Successful mating was confirmed by the presence of a vaginal plug the following morning, and noon of that day was designated as E0.5. Pregnant mice were then euthanized by cervical dislocation. Zygotes and 2-cell embryos were collected at E0.5 and E1.5, respectively, from the swollen ampullae of the oviducts (zygotes) or the oviducts (2-cell embryos) in prewarmed M2 medium (Sigma-Aldrich, MR-015-D) at E0.5 and E1.5, respectively

### Preparation of rat serum

To isolate rat serum (RS), 8-to 10-week-old male SD rats (∼300 g) were purchased from Beijing Sibefu Biotechnology Co., Ltd. Rats were anesthetized by intraperitoneal tribromoethanol (1.25% (v/v), 10 mL/kg; Nanjing Aibei Biotechnology Co., Ltd., M2960). Adequate anesthesia was confirmed by the absence of tail or hind paw reflex before exsanguination. Rats were placed in the supine position with their limbs secured, and the abdomen was sterilized with 75% (v/v) ethanol. The peritoneal cavity was opened, and the intestines were gently moved aside to expose the abdominal aorta. Blood was collected into anticoagulant-free vacuum tubes (Hebei Kangweishi Medical Technology Co., Ltd.), placed on ice to clot, and centrifuged at 2,500 × g for 10 min at 4°C. The serum was collected and centrifuged to remove any remaining red blood cells. The resulting serum was passed through a 0.22 μm filter, inactivated at 56°C for 30 min, divided into 1 mL aliquots, and stored at −80°C.

### Isolation of human umbilical cord blood serum

Immediately after birth, the umbilical cord was clamped 5-7 cm from the umbilicus and transected between the clamps. Umbilical cord blood was collected directly from the umbilical vein using a 14-gauge needle attached to a 50-mL syringe and transferred into sterile 15-mL centrifuge tubes without anticoagulants. The blood was allowed to clot at 4 °C and then centrifuged at 2,500 × g for 10 min at 4 °C. The serum was carefully collected and transferred to sterile centrifuge tubes. The isolated human umbilical cord blood serum (HCS) was stored at −80 °C until use. Before use, HCS was heat-inactivated in a water bath at 56 °C for 30 min, passed through a 0.22-μm filter, aliquoted, and stored at −80 °C. Repeated freeze-thaw cycles were avoided.

### Pre-implantation culture of mouse embryos

Collected zygotes were briefly treated with 0.2% (v/v) hyaluronidase (MedChemExpress, HY-107910) to remove granulosa cells. Zygotes or 2-cell embryos were washed three times in pre-equilibrated M2 medium and two times in pre-equilibrated KSOM (Sigma-Aldrich, MR-106-D) and cultured in 35-45 µL KSOM microdroplets under mineral oil (Vitrolife, 10029) at 37°C in 5% CO_2_ for 2 days (2-cell embryos) or 3 days (zygotes) s. Then, compacted morulae and early blastocysts were transferred to KSOM supplemented with 1× NEAA (Thermo Fisher Scientific, 11140050), and 1 mM sodium pyruvate (Gibco, 11360070) (designated KSOM+) and cultured until they reached the expanded or hatched blastocyst stage.

### Components of embryo culture medium

Extended *In Vitro* Culture medium 1 (EIVC1) consisted of CMRL1066 (Gibco, 1153037) supplemented with 20% heat-inactivated FBS, 1× NEAA, 1 mM sodium pyruvate, 1× GlutaMAX (Thermo Fisher Scientific, 35050061), 100 units/mL penicillin and 100 μg/mL streptomycin (Gibco, 15140122), 1× Insulin-Transferrin-Selenium-Ethanolamine (ITS-X) (Gibco, 51500056), 8 nM β-estradiol (Sigma-Aldrich, E8875), 200 ng/mL progesterone (Sigma-Aldrich, P0130), and 25 µM N-acetyl-L-cysteine (Sigma-Aldrich, A7250). EIVC2 consisted of CMRL1066 with 30% or 20% Knockout Serum Replacement (KSR) (Thermo Fisher Scientific, 10828028), 1× GlutaMAX, penicillin/streptomycin, and 1× ITS-X. For LDN treatment, LDN-193189 (MedChemExpress, HY-12071) was included as a component of the culture system. Modified *ex utero* culture medium (MEUCM) consisted of DMEM (Gibco, 11054020) supplemented with 1 mM sodium pyruvate, 1× GlutaMAX, 100 units/mL penicillin and 100 μg/mL streptomycin, and gradually increasing concentrations of heat-inactivated RS and human serum albumin (HSA) (Guangdong Shuanglin Bio-Pharmacy Co., Ltd).

### Post-implantation culture of mouse embryos

Expanded blastocysts were treated with EmbryoMax® Acid Tyrode’s solution (Sigma-Aldrich, MR-004-D) to remove the zona pellucida and washed at least three times with pre-equilibrated EIVC1 medium. Three to five zona-free or hatched blastocysts were then seeded into each well of a μ-Slide 8-well plate (Ibidi, 80826), designated as E0.5+D4 (zygotes) or E1.5+D3 (2-cell embryos). Before seeding, the wells were coated with 100 μL undiluted Matrigel (Corning, 354277) at 37 °C for approximately 30 min, followed by the addition of 350-400 μL pre-equilibrated EIVC1 containing 4% (v/v) Matrigel. After 24 h, the medium was replaced with 300 µL fresh EIVC1. After an additional 24 h, the medium was replaced with EIVC2 containing 30% KSR (Thermo Fisher Scientific, 10828028).

Embryos with abnormal morphology were discarded, and the medium was replaced with MEUCM. From E1.5+D7 or E0.5+D8 onward, MEUCM was refreshed daily using stage specific formulations: 10% RS, 10% KSR, and 1% HSA at E1.5+D7/E0.5+D8, and 30% RS with 3% HSA (without KSR) at E1.5+D9/E0.5+D10. At E1.5+D10, embryos that had developed to an E8.0-equivalent stage were transferred to roller culture bottles (2-4 embryos per bottle) containing 2-2.5 mL MEUCM with 30% RS and 3% HSA. The roller system was operated at 20-30 rpm and 37°C under an atmosphere of 5% O_2_, 21% CO_2_, and 74% N_2_ at 6.5 psi. The same culture conditions were maintained at E1.5+D11/D12 and E0.5+D12. For the optimized culture of zygote-derived embryos, HSA was replaced with HCS.

### Developmental stage alignment of cultured embryos

To align developmental stages between cultured and *in vivo* embryos, cultured embryos with similar developmental states were grouped according to their morphological correspondence to *in vivo* embryonic stages. Specifically, embryos from the E1.5+D9 and E1.5+D10 groups that morphologically corresponded to *in vivo* E8.0 were combined into the E1.5+D9/10 stage. Embryos from the E1.5+D10 and E1.5+D11 groups that morphologically corresponded to *in vivo* E8.5 were combined into the E1.5+D10/11 stage. Similarly, embryos from the E1.5+D11 and E1.5+D12 groups that morphologically corresponded to *in vivo* E9.0 were combined into the E1.5+D11/12 stage.

### Whole-mount immunostaining of *in vivo* and cultured mouse embryos

*In vivo* embryos at E4.75-E7.25 were dissected from pregnant mice, and the ectoplacental cone (EPC) and Reichert’s membrane were removed. For cultured embryos, only the EPC was removed, as these embryos lack Reichert’s membrane. For E8.5-E9.5 *in vivo* embryos and their corresponding cultured counterparts, the extraembryonic membranes (amnion, yolk sac, chorion, and allantois) were removed during dissection. After a brief wash with PBS (VivaCell, C3580-0500) at room temperature (RT; ∼26 °C), embryos were fixed in 4% paraformaldehyde (PFA; biosharp, BL539A) at 4 °C for 14-16 h. Embryos were then washed three times with PBS for 5 min each and stored in PBS until further use.

Embryos were subsequently permeabilized in PBS containing 0.5% Triton X-100 (Solarbio, T8200) and 0.1 M glycine (Millipore, 4810-OP) for 1-2 h at RT. After permeabilization, embryos were blocked in PBS containing 10% (v/v) FBS (Gibco, A5256701), 3% (m/v) bovine serum albumin (BSA) (Sigma-Aldrich, A2064), and 0.1% (v/v) Triton X-100. E4.75-E7.25 *in vivo* embryos and their cultured counterparts were blocked for 1-2 h at RT, whereas E8.0-E9.0 *in vivo* embryos and their corresponding cultured embryos were blocked for 16-18 h at 4 °C. Embryos were then incubated with primary antibodies diluted in blocking solution at 4 °C for 14-16 h. After three washes with PBS containing 0.1% Tween-20 (PBST; Sigma, P9416) for 5 min each, followed by two washes with PBS for 5 min each, embryos were incubated with corresponding secondary antibodies and DAPI (1:400; Sigma-Aldrich, D9542) in blocking solution for 2 h at RT. Finally, embryos were washed with PBST and stored in PBS at 4 °C until image acquisition. All primary and secondary antibodies used in this study are listed in Supplementary Table 6.

### Image acquisition, processing and analysis

Embryos were washed in PBS and mounted in SlowFade Glass Soft-Set Antifade Mountant (Thermo Fisher Scientific, S36917) between two No. 1.5 coverslips (Ibidi, NC1180546) with an appropriate spacer. Images were acquired using either a Stellaris 5 (Leica) or a Nikon AX laser scanning confocal microscope. For the Stellaris 5 system, a 16× HC Fluotar L 0.60 NA multi-immersion objective (Leica) and LAS X Navigator software (Leica) were used. For the Nikon AX system, the corresponding objective and software were applied according to the manufacturer’s recommendations. The following laser lines and emission detection windows were used: Hoechst (excitation 405 nm; emission 410-483 nm), Alexa Fluor 488 (excitation 488 nm; emission 493-556 nm), Alexa Fluor 568 (excitation 561 nm; emission 566-635 nm), and Alexa Fluor 647 (excitation 638 nm; emission 647-700 nm). For the Nikon AX system, 3D immunofluorescence (IF) analysis and visualization were used with IMARIS 10.1 (Oxford Instruments). Imaging parameters, including laser power, pixel dwell time, and detector gain, were carefully optimized to avoid photobleaching and signal saturation.

### IF staining of frozen sections of *in vivo* and cultured embryos

Fixed E9.0 embryos and their corresponding cultured counterparts were embedded in OCT compound (Sakura, 4583) and frozen at -20°C for at least 2 h. Embryos were then sectioned along the anterior-posterior (A-P) or dorsal-ventral (D-V) axis at a thickness of 15 μm using a cryostat (Leica, CM1950, Germany), and the sections were mounted onto pre-treated glass slides (CITOTEST, 1A5105). Frozen sections were stored at -80°C until IF staining. For IF staining, sections were equilibrated to RT for 30 min and washed three times with PBS. Antigen retrieval was performed by incubating the sections in HistoVT One solution (Nacalai Tesque, 06380-05) diluted 1:10 in ddH_2_O, in a water bath at 70°C for 20 min, followed by cooling to RT. Sections were permeabilized in PBS containing 0.3% Triton X-100 and 0.1 M glycine for 30 min and then blocked in PBS containing 10% FBS, 3% BSA, and 0.1% Triton X-100 at 4°C for 16-18 h. After blocking, sections were incubated with primary antibodies diluted in blocking solution at 4°C for 16-18 h, washed three times with PBST and three times with PBS, and then incubated with corresponding secondary antibodies and DAPI (1:400) for 2 h at RT. After three washes with PBST followed by three washes with PBS, sections were mounted and imaged using a Nikon AX laser confocal microscope (Nikon). All primary and secondary antibodies used in this study are listed in Supplementary Table 6.

### RNA extraction and quantitative real-time PCR

Approximately 15-20 embryos per group (E6.5 *in vivo* and E1.5+D7 *in vitro*) were collected. Total RNA was extracted using the RNeasy Mini Kit (Qiagen, Cat. 74004) and reverse-transcribed into cDNA using PrimeScript RT Master Mix (Takara, RR047A). Quantitative real-time PCR was performed using SYBR Green and gene-specific primers to measure *Id1* and *Id2* expression, with *Gapdh* as the internal control. Relative gene expression was calculated via the 2-ΔΔCt method, and data were analyzed using GraphPad Prism 8 (La Joya, California). Primer sequences used for quantitative real-time PCR are listed in the Supplementary Table 7.

### Cell preparation for scRNA-seq and scATAC-seq

Whole embryos were collected, rinsed in PBS to remove residual culture medium, and transferred to a digestion solution consisting of 1 mg/mL collagenase IV (Gibco, 17104019) and 0.25% trypsin-EDTA (Gibco, 25200072) mixed at a 1:1 ratio and supplemented with 100 ng/mL DNase I (Sigma-Aldrich, 10104159001). Embryos were gently shaken in a 37°C water bath for 1-3 min (depending on embryo size) and dissociated by gentle pipetting. Digestion was neutralized with PBS containing 10% FBS (Gibco, A5256701), and the resulting cell suspension was centrifuged at 300 × g for 3 min. The supernatant was removed, and the cells were resuspended in PBS containing 0.1% BSA (Sigma-Aldrich, B2064).

### Single-cell RNA-seq library preparation and sequencing

Cell viability was first assessed using a CountStar instrument (Countstar Rigel S2), and only cells with > 80% viability were further processed according to the 10× Genomics recommended protocol. Briefly, single cells were resuspended in 15-20 μL PBS containing 0.1% BSA at a concentration of 1,600-1,800 cells/µL for the Chromium GEM-X Single Cell 3’ v4 Chip (10× Genomics, PN-1000690) loading. Approximately 10,000–35,000 cells were loaded per channel, depending on sample conditions, resulting in the recovery of approximately 7,000–20,000 cells per channel. Then, single-cell transcriptome libraries were constructed using the Chromium GEM-X Single Cell 3’ Kit v4 (10× Genomics, PN-1000691). cDNA and library quality were assessed using a Qseq-100 instrument (BiOptic Inc), and qualified libraries were sequenced on an Illumina NovaSeq 6000 platform (Biolinker).

### Single-cell ATAC-seq library preparation and sequencing

Cell numbers were first determined using a cell-counting chamber (Countstar, CO010101). For samples containing fewer than 10,000 cells, the low-cell-input nuclei isolation protocol was followed. Wash buffer, lysis buffer, and diluted nuclei buffer were prepared on ice according to the 10× Genomics protocol (CG000169, Rev E). Fresh single-cell suspension was washed twice with 500 μL PBS containing 0.1% BSA and centrifuged at 400 × g for 5 min. After removal of the supernatant, cells were resuspended in 100 μL of lysis buffer and incubated on ice for 5 min to isolate nuclei. Subsequently, 1 mL wash buffer was added, and the nuclei were centrifuged at 500 × g for 5 min. The supernatant was removed, and the nuclei pellet was resuspended in diluted nuclei buffer and adjusted to a concentration of 1,200–5,500 nuclei/μL.

The prepared nuclei suspension was subjected to library construction in accordance with the instructions of the Chromium Next GEM Single Cell ATAC Kit v2 (10× Genomics, PN-1000390). Briefly, the nuclear suspension after transposition incubation was loaded into a 10× GEM Chip H (10× Genomics, PN-1000161), and then the nuclei were partitioned into nanoliter-scale Gel Beads in-emulsion (GEMs) suspensions through the Chromium Controller (10× Genomics). Subsequently, DNA amplification, recovery and library construction were carried out and the ATAC library quality was assessed by Qseq-100 instrument (BiOptic Inc). Finally, the library was sequenced on an Illumina NovaSeq 6000 platform by Biolinker.

### Spatial transcriptome sequencing of mouse embryos

E9.0 *in vivo* and E1.5+D11/D12 cultured embryos were collected, washed in cold PBS (4°C), and carefully dissected to remove the amnion, yolk sac, chorion, and allantois. Embryos were embedded in OCT compound, frozen on dry ice, and stored at -80°C until sectioning. Frozen embryos were sectioned at a thickness of 10 μm using a cryostat, mounted onto glass slides, stained with hematoxylin and eosin, and imaged with a microscope (Leica DM6B) to identify target regions. Adjacent sections were then mounted onto 1 cm × 1 cm Stereo-seq chips (STOmics, 200CT13114) and subjected to spatial transcriptome sequencing by Annoroad.

Briefly, chips were pre-incubated at 37°C for 5 min, fixed in pre-chilled methanol (Sigma-Aldrich, 34860) at -20°C for 30 min, stained with Qubit ssDNA dye (Thermo fisher Scientific, Q10212), and imaged with a Motic Custom PA53 FS6 microscope to assess tissue integrity. Sections were permeabilized with permeabilization solution at 37°C for 10-15 min, and the chips were washed with PR Rinse Buffer (STOmics, 1000033684) to release RNA. Reverse transcription was performed at 42°C for 5 h, followed by the addition of 400 μL tissue removal buffer and incubation at 55°C for 10 min. The chips were then washed with 0.1× SSC to remove residual tissue, and cDNA was released by incubating with 400 μL cDNA release buffer for 3 h. The reaction solution was collected, and cDNAs were purified using Agencourt AMPure XP beads (Agencourt, A63882). Purified cDNAs were amplified by PCR under the following conditions: initial denaturation at 95°C for 5 min; 15 cycles of 98°C for 20 s, 58°C for 20 s, and 72°C for 3 min; and a final extension at 72°C for 5 min. PCR products were purified with AMPure XP beads, and cDNA concentrations were measured using a Qubit fluorometer (Thermo Fisher Scientific, Q32854). For library construction, 20 ng of cDNA per sample was fragmented with Tn5 transposase (STOmics, 1000028515) at 55°C for 10 min, and the reaction was terminated with 5 μL of 0.02% SDS. A 75 μL PCR mix containing barcoded primers was then added, and amplification was performed under the following conditions: 95°C for 5 min; 13 cycles of 98°C for 20s, 58°C for 20s, and 72°C for 30s; and a final extension at 72°C for 5 min. The amplified products were purified with AMPure XP beads to generate a DNA nanoball (DNB) library, which was sequenced on a DNBSEQ™T7 platform with read lengths of 50 bp (read1) and 100 bp (read2).

### scRNA-seq data preprocessing, quality control, and integration

Raw BCL files were demultiplexed and converted to FASTQ format using bcl2fastq v2.20. Sequencing reads were then processed using Cell Ranger v9.0 (10× Genomics) with default parameters and aligned to the mouse reference genome (mm39).

The resulting gene expression count matrices were imported into Seurat v5.0.3^57^ for downstream analysis. Low-quality cells and potential artifacts were removed through multiple filtering steps. Potential doublets were identified and removed using DoubletFinder v2.0.4^58^ with default settings. Ambient RNA-contaminated cells were further identified and removed using the “decontX” method implemented in celda (v1.18.2)^59^. Cells were further filtered to retain those with 1,400-7,500 detected genes and a mitochondrial gene percentage of less than 5%. Filtered data were integrated using the FastMNNIntegration method implemented in SeuratWrappers v0.4.0^60^. Uniform manifold approximation and projection (UMAP) was performed using the first 30 principal components derived from the integrated data to generate a low-dimensional embedding for visualization. Cell clustering was performed using the “FindClusters” function in Seurat with a resolution of 1.

### Cell annotation and differentially expressed gene identification

Differentially expressed genes were identified using the “FindAllMarkers” function in Seurat with the Wilcoxon rank-sum test, and genes with an adjusted *P* value < 0.05 were retained as cluster-specific markers. Cell types were then annotated based on previously reported canonical marker genes. The same approach was applied for subcluster identification and annotation.

### ScATAC-seq quality control and integration

Raw scATAC-seq sequencing data were processed using the Cell Ranger ATAC pipeline (v2.1.0, 10x Genomics) and aligned to the mouse reference genome (mm39). The resulting fragment files and associated output files were used for downstream analyses. Quality control was performed using ArchR (v1.3.0)^61^. Cells with a transcription start site enrichment score > 7 and unique fragments > 1,000 were retained for downstream analyses. Potential doublets were identified and removed using the “filterDoublets” function in ArchR. Filtered scATAC-seq data were then subjected to dimensionality reduction and integration using ArchR. Iterative latent semantic indexing (Iterative LSI) was performed on the TileMatrix using three iterations and 20,000 variable features. Batch effects across samples were corrected using Harmony^62^ based on the Iterative LSI reduced dimensions, with “sample” as the integration variable. The Harmony-corrected low-dimensional embedding was used for clustering and visualization. Cell clusters were identified using the “addClusters” function in ArchR and preliminary peaks were called using the “addReproduciblePeakSet” function.

### ScATAC-seq cell-type annotation

The preliminary peak dataset was processed using Signac (v1.13.0)^63^ and Seurat. First, the PeakMatrix was normalized using term frequency-inverse document frequency (TF-IDF). Highly informative features were selected using the “FindTopFeatures” function, followed by singular value decomposition (SVD) for latent semantic indexing (LSI)-based dimensionality reduction via “RunSVD”. As the first LSI dimension is commonly associated with technical factors such as sequencing depth, 2nd-30th dimensions were retained for downstreamanalyses. To correct for batch effects across samples, Harmony integration was performed on the LSI embeddings, with “Sample” as the batch variable and project.dim = FALSE. UMAP visualization was performed using Harmony-corrected 2nd-30th dimensions. The same dimensions were used to construct the cell nearest-neighbor graph with the “FindNeighbors” function, followed by graph-based clustering with “FindClusters” in Seurat. Clusters with a median log10-transformed fragment count below 3.5 were excluded from downstream analyses. Cells were first assigned to major developmental lineages based on gene activity scores of canonical marker genes. Subsequently, constrained label transfer was performed separately for each lineage to assign refined cell-type identities based on the scRNA-seq dataset. Peaks were called for the resulting cell types using the “addReproduciblePeakSet” function in ArchR, generating a consensus peak set for downstream analyses.

### Construction of the TF regulatory network

SCENIC+(v 1.0a2)^31^ was applied to infer lineage-specific gene regulatory networks in cultured and *in vivo* embryos. Region sets used as input for SCENIC+ were constructed from the 3,000 NTOP topics, OTSU topics, and differentially accessible regions between lineages and between conditions (cultured and *in vitro*) within individual lineages. A custom cisTarget motif database was then generated with 1,000-bp background padding for each candidate regulatory region. Motif annotation was performed using motifs-v10nr_clust-nr.mgi as input. Finally, the generated region sets and custom cisTarget motif database were processed with SCENIC+ using default parameters to obtain enhancer-driven transcription factor-target gene modules, namely eRegulons.

### Stereo-seq quality control and data processing

Raw Stereo-seq data were processed using the SAW (v 8.1.3)^64^ pipeline for image registration, tissue segmentation, read alignment, and generation of spatial gene expression matrices. The resulting expression matrices were aggregated at bin50 resolution and imported into stereopy (v1.6.1)^65^ for downstream analysis. Low-quality spatial bins with fewer than 100 detected genes or a mitochondrial gene percentage greater than 5% were removed.

### Stereo-seq cell-type annotation

The filtered bin50 expression matrix was processed with stereopy for normalization and log-transformation. Highly variable genes were identified and used for Principal Component Analysis (PCA)-based dimensionality reduction, with the first 30 principal components retained for downstream analysis. Based on the PCA representation, a transcriptome-based nearest-neighbor graph was constructed using the “neighbors” function in stereopy. Spatial neighborhood relationships were then calculated using the “spatial_neighbors” function, based on the spatial coordinates of bin50 spots. The resulting spatial neighbor graph was used for UMAP visualization and Leiden clustering. The resulting clusters were used for spatial domain identification and annotation. Cell type identification was performed based on known marker gene sets.

### Anterior-posterior axis extraction and straightening

The embryonic anterior–posterior (A-P) axis was mapped, and the spatially curved embryo was computationally straightened along this axis for downstream analysis. Briefly, a series of control points were manually placed along the embryonic midline from the anterior to the posterior end to define the putative A-P centerline. A smooth spline curve was then fitted through these points to generate a continuous centerline. Each spatial point was assigned to its nearest position on the fitted centerline, and the arc length from the anterior starting point to that position was used as the coordinate along the straightened A-P axis. The perpendicular offset of each point from the centerline was also calculated to represent its relative position on either side of the axis. Thus, each original spatial point was transformed into a new two-dimensional coordinate system defined by A-P arc length and perpendicular distance.

### A-P axis profiling of ligand expression and cell type distribution

Cells were projected onto the straightened A-P coordinate system, and the coordinate along this axis was used as the A-P score. Cells ordered by A-P score were assigned to sliding windows, each spanning 2% of the total A-P score range. For each window, the aggregated log-transformed expression of ligand genes was calculated and normalized by the number of cells within the window, and cell type composition was determined as the fraction of cells assigned to each type. Spatial regions along the A-P axis that exhibited both elevated ligand expression and enrichment of specific cell types were defined as potential signaling centers. Ligand, receptor, activator, and inhibitor gene sets were extracted from all signaling pathways in the CellChat database (CellChatDB.mouse)^35^.

### Cell-cell communication analysis

Cell type annotations and normalized expression matrices were extracted from Seurat objects and used as input for CellChat (v 2.2.0)^35^ analysis using the mouse ligand-receptor interaction database (CellChatDB.mouse). Cell-cell communication inference was performed following the standard CellChat workflow. Communication probabilities were calculated using the “computeCommunProb” function, and weak interactions were removed using the “filterCommunication” function with default settings. Pathway-level communication probabilities were computed using the “computeCommunProbPathway” function, and aggregated communication networks were generated using the “aggregateNet” function. Network centrality was calculated using the “netAnalysis_computeCentrality” function, and incoming and outgoing signaling patterns were visualized using the “netAnalysis_signalingRole_heatmap” function.

### Functional enrichment analysis

Gene Ontology (GO) enrichment analysis and gene set enrichment analysis (GSEA) were performed using the “enrichGO” and “GSEA” functions, respectively, in the clusterProfiler package (v4.10.1)^66^.

### Single-cell transcriptomic comparison of *in vivo* and cultured mouse embryos

Label transfer analysis was performed using the scRNA-seq dataset of cultured E1.5+D11/12 embryos as the query and the E8.5, E8.75, and E9.0 scRNA-seq datasets^17^ as the references. Cell types from both *in vivo* and *in vitro* embryos were first grouped into seven major lineages. Transfer anchors were identified using the “FindTransferAnchors” function in Seurat based on the first 30 principal components, followed by lineage label transfer using “TransferData”. The proportions of *in vitro* cells assigned to each *in viv*o developmental stage-associated lineage were then calculated to assess developmental correspondence between the two conditions.

### Embryos collection and metabolite extraction

Embryos at specific developmental stages were collected in biological triplicates. Fresh embryos were rinsed three times in 3 mL of ice-cold PBS containing 0.25% polyvinylpyrrolidone (PVP; Sigma-Aldrich, PVP40) and subsequently flash-frozen in liquid nitrogen^67^. All samples were stored in liquid nitrogen until metabolite extraction. For metabolite extraction, frozen embryo pools were transferred to dry ice-chilled tube containing 100 µL of extraction buffer (methanol (Thermo Scientific Chemicals, A456-4): acetonitrile (Thermo Scientific Chemicals, A955-4): ddH_2_O, 40:40:20, v/v/v). Embryos were mechanically homogenized using an electronic tissue grinder (TIANGEN, OSE-Y50) at 11,000 rpm in 3s pulses with 3s pauses for a total of 1 min, while ensuring that the samples remained frozen. The homogenate was then sonicated on a pre-chilled metal rack using a pulse program (3s on, 3s off) for 1 min. The lysate was centrifuged at 15,000 × g for 30 min at 4°C. The resulting supernatant was transferred to a new 1.5mL Eppendorf tube and flash-frozen, while the pellet was reserved for protein quantification via a bicinchoninic acid (BCA) assay (NCM Biotech, WB6501) after lysis in RIPA buffer (Beyotime, P0013B). Protein concentration was used for subsequent data normalization. The frozen supernatant was dried under vacuum at -20°C for 1 h. The dried metabolites were reconstituted in 80% (v/v) acetonitrile on ice and centrifuged again at 15,000 × g for 30 min at 4°C to remove insoluble debris.

### LC-MS/MS data acquisition

Samples were reconstituted in 40 µL of 80% acetonitrile, and a 5-µL aliquot of the supernatant was injected for LC-MS analysis. Chromatographic separation was performed using a Vanquish UHPLC system (Thermo Fisher Scientific) equipped with a Waters ACQUITY UPLC BEH Amide column (2.1 × 100 mm, 1.7 µm). The column temperature was maintained at 40°C, and the flow rate was set to 0.5 mL/min. The mobile phases consisted of 25 mM ammonium acetate (Sigma-Aldrich, 5.33004) and 25 mM ammonium hydroxide (Solstice, 60-046-886) in ddH_2_O (pH 9.75) (A) and acetonitrile (B) (Thermo Scientific Chemicals, A955-4). The gradient elution program was as follows: 95% B from 0 to 0.5 min; 95-65% B from 0.5 to 7 min; 65-40% B from 7 to 8 min; 40% B from 8 to 9 min; 40-95% B from 9 to 9.1 min; and 95% B from 9.1 to 12 min.

Mass spectrometric detection was performed on a Q Exactive HF-X mass spectrometer (Thermo Fisher Scientific) equipped with a heated electrospray ionization (HESI) source and operated in both positive and negative ionization modes. Untargeted MS/MS data were acquired using a normalized collision energy (NCE) of 30. For targeted analysis, compounds were identified by matching accurate mass (m/z) and retention time to authentic standards. Targeted data analysis was conducted using Thermo Xcalibur software (v4.2). Sample information is provided in Supplementary Table 8.

### Metabolic data analysis

Quality control (QC) was injected three times before sample sequences and every 9 samples, with good repeatability after the first two injections. 78.32% of features in positive ion mode and 72.19% in negative ion mode passed the 30% relative standard deviation (RSD) cutoff, indicating good-quality metabolomics data for subsequent analysis using the tidymass framework (https://www.tidymass.org) built in R (v.4.5.2)^68^. Only features present in at least 80% of QC samples were retained. After blank subtractions, the sample data were normalized to each sample’s protein concentration by the BCA. No batch effect was assumed. No outlier was detected. The missing values were imputed by the k-nearest neighbors (KNN) algorithm with default parameters. Then, median normalization was performed within the tidymass pipeline. Metabolite annotation was performed using the metID package (v.1.31.0), which utilizes MS1 m/z, retention time, and MS/MS spectral matching against three databases (Human Metabolome Database (HMDB), MassBank of North America (MoNA), Massbank) with default parameters and the column type set to hydrophilic liquid chromatography (HILIC). A two-tier annotation confidence was adopted, following the Metabolomics Standards Initiative guidelines, and the top-scoring annotation was retained when multiple annotations were present. Sarcosine was assigned to M88T326_NEG instead of alanine based on targeted data. Then, the z-score standardized abundances of annotated metabolites from tidymass metabolomics were pooled with the z-score standardized abundances of targeted metabolites quantified using in-house standards. For certain metabolites detected in both platforms (e.g., lactate), the strong concordance between datasets validated the reliability of our untargeted metabolomics processing workflow. For these overlapping metabolites, we prioritized the targeted measurements and then selected the features with the minimal standard deviation across biological replicates. In total, 261 metabolites were retained for subsequent analysis. Next, the ClusterGVis package^69^ was customized to enable cross-group temporal visualization of metabolite clusters. Pathway enrichment analysis was conducted using MetaboAnalystR (v.4.2.0)^70^ against the Small Molecule Pathway Database (SMPDB)^71^, and only statistically significant results (P < 0.05) were visualized in the figures.

### Statistical analysis

All statistical analyses were performed using GraphPad Prism 8 software (La Joya, California). Metabolomics data are presented as mean ± s.e.m. from a minimum of two independent experiments, whereas embryonic data are presented as mean ± S.D. A Kolmogorov–Smirnov test was performed to check normal distribution of data before each statistical test. Significant differences between two samples were evaluated using an unpaired two-sided Student’s t-test for normally distributed data or a Mann–Whitney test for non-normally distributed data. P < 0.05 was considered as statistically significant.

## Acknowledgements

This work was funded by grants from the National Natural Science Foundation of China (82425025, 32288101, 82225019, 82192871, and 32300687), the National Key Research and Development Program of China (2023YFA1801802, 2024YFA1106900, 2024YFC2706600, 2023YFA1800300, and 2025YFA1804400), the Major Basic Research Project of Science and Technology of Yunnan (202401BC070020, 202601AS070083, and 202401AT070097) and the Collaborative Research Fund grant C4007-24E to C.S..

## Author contributions

T.T. and J.W. conceptualized this project. T.T., Y.Y., B.B., and C.S. supervised the overall experiments. H.Y., N.S., Y.F., H.S., and J.H. performed embryo culture experiments. T.X. H.Y., N.S., H.S., and J.H. performed the metabolic experiments. T.X. performed the metabolic data analysis. H.Y., N.S., H.S., Y.Z., and L.F. isolated embryonic single cells and subsequently constructed scRNA-seq, scATAC-seq, and stereo-seq libraries. K.W., B.C., and B.B. performed all the bioinformatic analysis. Y.L., Y.D., T.X., and Z.Y. provided technical support in the experiments. J.W. offered feedback and reviewed the manuscript. T. T., J.W., B.B., and N.S. prepared the manuscript with input from the other authors.

## Additional information

Correspondence and requests for materials should be addressed to Weizhi Ji, Chun So, Bing Bai, Yang Yu, Jun Wu, or Tao Tan.

