## Extended data figure legends for "Continuous *ex utero* development of mouse zygotes to early organogenesis"

**Extended Data Fig. 1 Optimization of the pre-implantation culture system for mouse embryos. a**, Schematic diagram of the optimization strategy for mouse pre-implantation embryo culture medium from E1.5 to D3 (Dx = E1.5+Dx). **b**, Representative bright-field images of embryos cultured in KSOM for the first two days (D1-D2) and then in either KSOM or KSOM+ for the third day (D2–D3), starting from E1.5. Scale bar: 200 μm. **c,** Quantification of blastocyst hatching ratio at D3 in the KSOM and KSOM+ groups. n = 187 (KSOM, 11 independent experiments) and n = 218 (KSOM+, 10 independent experiments). Data are shown as mean ± S.D. *****P* < 0.0001 (unpaired t-test). Representative bright-field images are shown below each graph. Scale bar, 100 μm. **d-e**, Quantification of blastocyst diameter in hatched (**d**) and unhatched (**e**) blastocysts at D3 in the KSOM and KSOM+ groups. The red dotted line indicates the measurement axis from the polar to the mural trophectoderm (TE). Hatched: n = 12 (KSOM) and n = 28 (KSOM+). Unhatched: n = 16 (KSOM) and n = 21 (KSOM+). Data are shown as mean ± S.D. ****P* < 0.001; ***P* < 0.01 (unpaired t-test). Representative images are shown below each graph. Scale bar, 100 μm. **f,** Quantification of embryos with lumpy mural TE (mTE) at D4 in the KSOM and KSOM+ groups. Red arrows indicate lumpy mTE. n = 112 (KSOM, 4 independent experiments) and n = 135 (KSOM+, 4 independent experiments). Scale bar, 100 μm. **g-h**, Representative bright-field images (**g**) and quantification (**h**) of egg cylinder embryo formation at D6 in the KSOM and KSOM+ groups. Red dotted circles delineate egg cylinder-shaped embryos. n = 112 (KSOM, 4 independent experiments) and n = 135 (KSOM+, 4 independent experiments). Data are shown as mean ± S.D. *****P* < 0.0001 (unpaired t-test). Scale bar, 200 μm.

**Extended Data Fig 2 Organogenesis failure in embryos cultured in the unoptimized system. a,** Representative bright-field images showing morphology of cultured mouse embryos from the 2-cell stage to D10 (Dx = E1.5+Dx). Scale bars, 100 μm (E1.5-D7) and 200 μm (D8-D10). **b,** Developmental ratio of cultured mouse embryos from D3 to D11. n = 188 embryos from 5 independent experiments. Data are shown as mean ± S.D. **c**, Schematic (left) and quantification (right) of EPI length from D6 to D8 and in *in vivo* counterparts from E5.5 to E7.25. Green double-headed arrows indicate the axis used for EPI length measurements. n = 14 (D6), 9 (D7), 10 (D8), 15 (E5.5), 14 (E6.5), and 8 (E7.25). Data are shown as mean ± S.D. ns, not significant (unpaired t-test). **d**, Representative whole-mount immunofluorescence (IF) images of D5 and E5.25 embryos stained for ZO1 (green), SOX2 (magenta), and DAPI (blue). n = 4 (D5) and n = 3 (E5.25). Scale bars, 100 μm. **e,** Representative whole-mount IF images of D6 and E5.5 embryos stained for CER1 (green), TOX3 (magenta), LEFTY1 (yellow), and DAPI (blue). n = 3 (D6) and n = 4 (E5.5). Scale bars, 50μm. **f,** Representative whole-mount IF images of D8 and E7.25 embryos stained for T (green), FOXA2 (magenta), and DAPI (blue). Lower panels show higher-magnification views of the boxed regions in the upper panels; white arrows indicate T⁺FOXA2⁺ axial mesoderm cells. n = 5 (D8) and n = 4 (E7.25). Scale bars, 200 μm. ICM, inner cell mass; TE, trophectoderm; EPI, epiblast; ExE, extraembryonic ectoderm; EPC, ectoplacental cone; Am, amnion; VE, visceral endoderm; DVE/ AVE, distal visceral endoderm/anterior visceral endoderm; ExMe, extraembryonic mesoderm; PGC, primordial germ cells.

**Extended Data Fig. 3 Abnormal activation of BMP signaling in mouse embryos cultured in the unoptimized system. a**, Three-dimensional UMAP visualization of integrated scRNA-seq datasets from D5-D8 (Dx = E1.5+Dx) embryos cultured in the unoptimized system (left) and publicly available *in vivo* E5.0-E7.25 datasets (right)^1,2^. Each color represents a distinct cell type. Cell types labeled in red indicate those absent in cultured embryos (non-neural ectoderm) or *in vivo* embryos (EPC). **b**, Dot plot of scaled marker gene expression across cell types in scRNA-seq data from D5-D8 embryos cultured in the unoptimized system. Bubble size indicates the proportion of cells expressing each gene; color intensity indicates expression level. **c**, UpSet plots showing shared GO terms related to signaling among upregulated (left) and downregulated (right) DEGs in unoptimized cultured embryos versus *in vivo* embryos (|log2FC| > 1, adjusted *P* < 0.05). Vertical bars indicate the intersection sizes of shared GO terms across cell types; horizontal bars indicate the total number of GO terms per cell type. Connected dots indicate the cell types included in each intersection. **d**, Gene set enrichment analysis (GSEA) of BMP, WNT, and TGF-β receptor (TGF-β R) signaling pathways across major cell types using upregulated DEGs between unoptimized cultured embryos (D5-D8) and their *in vivo* counterparts. The color scale indicates column-scaled normalized enrichment scores (NES) across cell types. **e**, Heatmap showing average expression of BMP pathway-related genes (*Id1*, *Id2*, *Bmpr1a*, *Bmpr1b*, *Bmpr2*, and *Acvr1*) across cell types in unoptimized cultured (D5-D8) and *in vivo* (E5.0-E7.25) embryos. **f**, Bar plot showing average expression of BMP pathway-related genes (*Id1*, *Id2*, *Bmpr1a*, *Bmpr1b*, *Bmpr2*, and *Acvr1*) in EPI cells from *in vivo* E5.5^1^, unoptimized cultured D6 embryos, and stem cell-derived embryo models^3,4^ at the E5.5-equivalent stage. Data are shown as mean ± S.D. Box colors indicate the dataset source. EPI, epiblast; PS, primitive streak; PGC, primordial germ cells; Nas.M, nascent mesoderm; LPM, lateral plate mesoderm; HEP, haemato-endothelial progenitors; ExE, extraembryonic ectoderm; VE, visceral endoderm; PE, parietal endoderm; ExEn, extraembryonic endoderm; EPC, ectoplacental cone.

**Extended Data Fig. 4 Molecular and morphological characterization of early organogenesis in cultured embryos after BMP inhibition. a,** Relative expression of *Id1* and *Id2* in E6.5, LDN-, and LDN+ whole embryos, as determined by qPCR. Data are shown as mean ± S.D.; n = 9 from independent experiments. **P* < 0.05, ***P* < 0.01, *****P* < 0.0001 (paired t-test). **b,** Heatmap showing average normalized expression of *Id1* and *Id2* across cell types in scRNA-seq data from *in vivo* E6.5^2^ and cultured embryos with or without LDN treatment. **c,** Representative bright-field images of LDN-treated embryos at D9 and D10, alongside stage-matched *in viv*o E8.0 and E8.5 embryos. Scale bar, 200 μm. **d,** Representative whole-mount IF images of cultured D10 and *in vivo* E8.5 embryos stained for DAPI (blue), SOX2 (yellow), and HOXB4 (green). Lower panels show higher-magnification views of the boxed regions in the upper panels, highlighting the somites and neural tube. n = 2 (D10) and n = 6 (E8.5). Scale bar, 500 μm. **e,** Representative whole-mount IF images of cultured D10 and *in vivo* E8.5 embryos stained for DAPI (blue), SOX2 (yellow), and MYH2 (magenta). Lower panels show higher-magnification views of the boxed regions in the upper panels; green dotted lines outline the heart tube. n = 3 (D10) and n = 5 (E8.5). Scale bars, 500 μm (upper panels) and 200 μm (lower panels). **f,** Representative whole-mount IF images of cultured D10 and *in vivo* E8.5 embryos stained for FOXG1 (green), OTX2 (yellow), and DAPI (blue). Lower panels show higher-magnification views of the boxed regions in the upper panels, highlighting forebrain and midbrain patterning. n = 2 (D10) and n = 2 (E8.5). Scale bars, 500 μm (upper panels) and 200 μm (lower panels). **g,** Representative whole-mount IF images of cultured D10 and *in vivo* E8.5 embryos stained for CDX2 (green), ISL1 (yellow), and DAPI (blue). Lower panels show higher-magnification views of the boxed regions in the upper panels. Yellow arrows indicate the heart tube; yellow arrowheads denote the foregut and hindgut. n = 2 (D10) and n = 3 (E8.5). Scale bars, 500 μm (upper panels) and 200 μm (lower panels). **h,** Representative whole-mount IF images of cultured D10 and *in vivo* E8.5 embryos stained for CDX2 (green), ISL1 (yellow), and DAPI (blue). Yellow arrows indicate the neural tube. n = 2 (D10) and n = 3 (E8.5). Scale bars, 500 μm. EPI, epiblst; PS, primitive streak; ExE, extraembryonic ectoderm; ExEn, extraembryonic endoderm; PE, parietal endoderm; VE, visceral endoderm; YS, yolk sac; EPC, ectoplacental cone; Al, allantois; Am, amnion; NT, neural tube; Hf, head fold; Fb, forebrain; Mb, midbrain; Hb, hindbrain; Fg, foregut; Ht, heart; So, somites; Tb, tail bud.

**Extended Data Fig. 5 Neural and cardiac development in cultured 2-cell embryos at D11/12. a,** Diagram (left) and representative bright-field images (right) showing axial rotation at D11/12 (Dx = E1.5+Dx). Embryos were classified into four stages: (1) completely unturned; (2) ~50% turned; (3) > 50% but incompletely turned; and (4) fully turned. n = 34 from 7 independent experiments. Scale bars, 400 μm. **b,** Representative IF images of transverse sections of the neural tube region in D11/12 and E9.0 embryos stained for SOX9 (green) and TWIST (yellow). Lower panels are magnified views of boxed regions; white arrows indicate SOX9⁺TWIST⁺ cells. n = 3 per stage. Scale bars, 50 μm. **c,** Representative IF images of transverse sections from D11/12 and E9.0 embryos stained for SOX2 (magenta), FOXA2 (green), and NKX2-2 (yellow). n = 4 (D11/12) and n = 3 (E9.0). Scale bars, 50μm. **d,** Representative whole-mount IF images of cultured D11/12 and E9.0 embryos stained for MYH2 (magenta), SOX2 (yellow), and DAPI (blue). Lower panels show magnified views of the boxed regions; green dotted lines outline the heart. n = 6 (D11/12) and n = 4 (E9.0). Scale bars, 500 μm (upper panels) and 200 μm (lower panels). **e,** Schematic of E9.0 mouse heart morphology and the corresponding section planes in D11/12 and E9.0 embryos. **f,** Representative IF images of transverse sections from D11/12 and E9.0 embryos stained for TBX5 (magenta) and ISL1 (yellow). Lower panels show magnified views of the boxed regions. n = 4 (D11/12) and n = 3 (E9.0). Scale bar, 100 μm.

**Extended Data Fig. 6 Cell type annotation and integration of *in vitro* and *in vivo* scRNA-seq and scATAC-seq datasets. a-c**, UMAP plots showing 51 major cell types in scRNA-seq data from cultured D5-D11/12 embryos (**a**) and scATAC-seq data from cultured D6-D11/12 embryos (**b**) and *in vivo* E5.5-E9.0 embryos (**c**). Each color represents a distinct cell type. **d**, Dot plot showing scaled expression of marker genes across cell types in scRNA-seq data from cultured D5-D11/12 embryos. Dot size represents the percentage of expressing cells; color intensity represents the expression level. **e**, Dot plot showing scaled gene activity scores of marker genes across cell types in scATAC-seq datasets from cultured D6-D11/12 and *in vivo* E5.5-E9.0 embryos. Dot size represents the percentage of active cells; color intensity represents the gene activity score. **f**, Heatmap showing the percentage of major cell types from *in vivo* embryos (E8.5-E9.0) mapped to cultured D11/12 cell types by label transfer. **g**, UMAP plot of *Nkx2-1*⁺ cells within the pharyngeal endoderm in scRNA-seq data from cultured D11/12 embryos (left) and dot plot showing normalized expression of lung progenitor marker genes in these cells (right). Dot size indicates the percentage of expressing cells; color intensity indicates the expression level. EPI, epiblast; PS, primitive streak; PGC, primordial germ cells; Cardiomyocytes SHF, second heart field-derived cardiomyocytes; Cardiomyocytes FHF, first heart field-derived cardiomyocytes; LPM, lateral plate mesoderm; YS mesothelium, yolk sac mesothelium; HEC, haemato-endothelial cells; EMP, erythroid-myeloid progenitors; NMP, neuro-mesodermal progenitors; FMH-brain, forebrain/midbrain/hindbrain; EPC, ectoplacental cone; ExE, extraembryonic ectoderm; ExEn, extraembryonic endoderm; Nas.M, nascent mesoderm; VE, visceral endoderm; PE, parietal endoderm; E, embryonic day; PrE, primitive endoderm.

**Extended Data Fig. 7 Spatial transcriptomic characterization of cell types and intercellular communication in cultured D11/12 and E9.0 embryos. a-b**, UMAP visualization of the cultured D11/12 (**a**) and *in vivo* E9.0 (**b**) Stereo-seq datasets. Cells are colored according to 19 annotated clusters. Dx = E1.5+Dx. **c-d**, Dot plots showing normalized and log1p-transformed expression of marker genes across cell types in D11/12 (**c**) and E9.0 (**d**) Stereo-seq datasets (bin size = 50), respectively. Dot size represents the percentage of expressing cells; color intensity represents the expression level. **e**, Spatial plots showing annotated cell type distribution across embryo sections in D11/12 and E9.0 Stereo-seq datasets. Colored dots indicate cell types. **f**, Heatmaps showing scaled average expression of CellChat-derived ligand genes across cell types in D11/12 and E9.0 Stereo-seq datasets. Color intensity from blue to red indicates low to high expression levels. **g,** Heatmaps showing outgoing (left) and incoming (right) signaling among cardiomyocytes, primitive gut, and pharyngeal mesoderm, derived from cultured D11/12 (top) and *in vivo* E9.0 (bottom) Stereo-seq datasets. Green color intensity indicates relative signaling strength. Colored bars above the heatmaps indicate the total signaling strength of each cell type, whereas gray bars on the right indicate the total signaling strength of each signaling pathway. **h**, Spatial plots showing expression of LAMININ pathway ligand and receptor genes in cultured D11/12 (left) and E9.0 (right) Stereo-seq datasets. Dashed boxes indicate putative interaction regions between the primitive gut (blue) and cardiomyocytes (red). LPM, lateral plate mesoderm; PLPM, posterior lateral plate mesoderm; FMH-brain, forebrain/midbrain/hindbrain; NMP, neuro-mesodermal progenitors.

**Extended Data Fig. 8 Quality control and annotation confidence of untargeted metabolomics data. a-d,** Quality control of untargeted metabolomics data. Total ion count (TIC) chromatograms over retention time for quality control samples (QC 1-8) in positive ion mode (POS, **a**) and negative ion mode (NEG, **c**). Accumulated percentage of detected features plotted against residual standard deviation (RSD) in POS (**b**) and NEG (**d**). Red vertical lines mark the 30% RSD threshold; numbers in red squares indicate the corresponding percentage of features retained at this threshold (78.32% for POS, 72.19% for NEG). **e,** Heatmap showing normalized abundance (Z-score) of metabolites detected by both targeted and untargeted metabolomics analyses. **f,** Pearson’s correlation coefficients between POS and NEG data for metabolites detected in both ionization modes. **g,** Pearson’s correlation coefficients between targeted and untargeted metabolomics data for metabolites detected by both methods. **h,** Mean standard deviation (SD) of Z-score-normalized abundance across biological replicates for each metabolite.

**Extended Data Fig. 9 Metabolic pathway enrichment and metabolite dynamics in Clusters 1 and 4 between *in vivo* and cultured embryos**. **a**, Network showing significantly enriched pathways (P < 0.05; blue circles) and their associated metabolites (yellow circles) in Cluster 1. **b**, Normalized abundance of creatine across developmental stages (*in vivo* E5.5-E9.0; *in vitro* D6-D12; Dx = E1.5+Dx), fitted with cubic polynomial regression (solid lines) with 95% confidence intervals (shaded areas). The interaction term (q_int) between developmental stage and culture group is significant (*P* < 0.01). **c**, Normalized abundance of fumarate across developmental stages (*in vivo* E5.5-E9.0; *in vitro* D6-D12), fitted with cubic polynomial regression (solid lines) and 95% confidence intervals (shaded areas). The interaction term (q_int) between stage and culture group is significant (*P* < 0.0001). **d**, Normalized abundance of taurine across developmental stages (*in vivo* E5.5-E9.0; *in vitro* D6-D12), fitted with cubic polynomial regression (solid lines) and 95% confidence intervals (shaded areas). The interaction term (q_int) between stage and culture group is significant (*P* < 0.05). **e**, Normalized abundance of propionylcarnitine across developmental stages (*in vivo* E5.5-E9.0; *in vitro* D6-D12), fitted with cubic polynomial regression (solid lines) and 95% confidence intervals (shaded areas). The interaction term (q_int) between stage and culture group is significant (*P*<0.01). **f**, Normalized abundance of butyrylcarnitine across developmental stages (*in vivo* E5.5-E9.0; *in vitro* D6-D12), fitted with cubic polynomial regression (solid lines) and 95% confidence intervals (shaded areas). The interaction term (q_int) between stage and culture group is significant (*P*<0.01). **g**, Normalized abundance of palmitoylcarnitine across developmental stages (*in vivo* E5.5-E9.0; *in vitro* D6-D12), fitted with cubic polynomial regression (solid lines) and 95% confidence intervals (shaded areas). The interaction term (q_int) between stage and culture group is significant *P*<0.01). **h**, Network showing significantly enriched pathways (P < 0.05; blue circles) and their associated metabolites (yellow circles) in Cluster 4. **i**, Normalized abundance of methionine across developmental stages (*in vivo* E5.5-E9.0; *in vitro* D6-D12), fitted with cubic polynomial regression (solid lines) and 95% confidence intervals (shaded areas). The interaction term (q_int) between stage and culture group is significant (*P* < 0.0001). **j**, Normalized abundance of oxoglutarate (α-KG) across developmental stages (*in vivo* E5.5-E9.0; *in vitro* D6-D12), fitted with cubic polynomial regression (solid lines) and 95% confidence intervals (shaded areas). The interaction term (q_int) between stage and culture group is significant (*P* < 0.0001). **k**, Normalized abundance of succinate across developmental stages (*in vivo* E5.5-E9.0; *in vitro* D6-D12), fitted with cubic polynomial regression (solid lines) and 95% confidence intervals (shaded areas). The interaction term (q_int) between stage and culture group is significant (*P* < 0.0001). **l**, Normalized abundance of glutamate across developmental stages (*in vivo* E5.5-E9.0; *in vitro* D6-D12), fitted with cubic polynomial regression (solid lines) and 95% confidence intervals (shaded areas). The interaction term (q_int) between stage and culture group is significant (*P* < 0.0001). **m**, Normalized abundance of glutamine across developmental stages (*in vivo* E5.5-E9.0; *in vitro* D6-D12), fitted with cubic polynomial regression (solid lines) and 95% confidence intervals (shaded areas). The interaction term (q_int) between stage and culture group is significant (*P* < 0.001). **n**, Normalized abundance of phosphate across developmental stages (*in vivo* E5.5-E9.0; *in vitro* D6-D12), fitted with cubic polynomial regression (solid lines) and 95% confidence intervals (shaded areas). The interaction term (q_int) between stage and culture group is significant (*P* < 0.01).

**Extended Data Fig. 10 Glycolytic pathway dynamics in metabolic and transcriptomic profiles of embryos cultured from the 2-cell stage. a**, Network showing significantly enriched pathways (blue circles) and their associated metabolites (yellow circles) in Cluster 5. **b**, Heatmap showing normalized average expression of glycolytic enzymes (white-purple-yellow scale; increasing expression) and normalized average abundance of glycolytic intermediates (blue-white-red scale; increasing abundance) across developmental stages (*in vivo* E5.5-E9.0; *in vitro* D6-D12; Dx = E1.5+Dx). **c**, Normalized abundance of glucose-6-phosphate across developmental stages (*in vivo* E5.5-E9.0; *in vitro* D6-D12), fitted with cubic polynomial regression (solid lines) and 95% confidence intervals (shaded areas). The interaction term (q_int) between stage and culture group is not significant (*P* > 0.05). **d**, Normalized abundance of fructose-6-phosphate across developmental stages (*in vivo* E5.5-E9.0; *in vitro* D6-D12), fitted with cubic polynomial regression (solid lines) and 95% confidence intervals (shaded areas). The interaction term (q_int) between stage and culture group is not significant (*P* > 0.05). **e**, Normalized abundance of dihydroxyacetone phosphate across developmental stages (*in vivo* E5.5-E9.0; *in vitro* D6-D12), fitted with cubic polynomial regression (solid lines) and 95% confidence intervals (shaded areas). The interaction term (q_int) between stage and culture group is not significant (*P* > 0.05). **f**, Normalized abundance of ribose-5-phosphate across developmental stages (*in vivo* E5.5-E9.0; *in vitro* D6-D12), fitted with cubic polynomial regression (solid lines) and 95% confidence intervals (shaded areas). The interaction term (q_int) between stage and culture group is not significant (*P* > 0.05). **g**, Normalized abundance of glycerol-3-phosphate across developmental stages (*in vivo* E5.5-E9.0; *in vitro* D6-D12), fitted with cubic polynomial regression (solid lines) and 95% confidence intervals (shaded areas). The interaction term (q_int) between stage and culture group is not significant (*P* > 0.05). **h**, Normalized abundance of lactate across developmental stages (*in vivo* E5.5-E9.0; *in vitro* D6-D12), fitted with cubic polynomial regression (solid lines) and 95% confidence intervals (shaded areas). The interaction term (q_int) between stage and culture group is not significant (*P* > 0.05).

**Extended Data Fig. 11 The 2-cell culture system supports zygote development *in vitro* for up to 12 days. a,** Developmental ratio of cultured mouse embryos from D4 to D12 (Dx = E0.5+x). n = 205 from 7 independent experiments. Data are shown as mean ± S.D. **b,** Representative whole-mount IF images of cultured D5 and D7 embryos and *in vivo* E4.75 and E5.5 embryos stained for CDX2 (green), OCT4 (magenta), GATA6 (yellow), and DAPI (blue). n = 2 (D5), 2 (D7), 5 (E4.75), 3 (E5.5). Scale bars, 100 μm. **c,** Representative whole-mount IF images of cultured D8 and *in vivo* E6.5 embryos stained for T (green), DKK1 (yellow), and DAPI (blue). n = 4 (D8) and n = 6 (E6.5). Scale bars, 100 μm. **d,** Representative whole-mount IF images of cultured D9 and *in vivo* E7.25 embryos stained for T (green), FOXA2 (magenta), OTX2 (yellow), and DAPI (blue). Upper panels show whole embryos; middle panels show single-layer magnified views of the boxed regions in the upper panels; lower panels show higher-magnification views of the boxed regions in the middle panels. Red arrowheads indicate FOXA2⁺OTX2⁺ visceral endoderm cells; white arrows denote OTX2⁺ prospective anterior neuroectoderm; white dotted circles outline organizer cells. n = 2 (D9) and n = 3 (E7.5). Scale bars, 200 μm (upper and middle) and 100 μm (lower). **e,** Representative bright-field images of cultured D12 (upper) and *in vivo* E8.75 (lower) embryos. Scale bar, 500 μm. **f,** Representative whole-mount IF images of cultured D12 and *in vivo* E8.75 embryos stained for SOX2 (yellow) and DAPI (blue). Somites are numbered, and their positions are indicated. n = 4 (D12) and n = 10 (E8.75). Scale bar, 500 μm. **g,** Quantification of somite pair numbers in cultured D12 and *in vivo* E8.75 embryos. n = 4 (D12) and 10 (E8.75). Data are shown as mean ± S.D. ns, not significant (unpaired t-test). **h,** Representative whole-mount IF image of a cultured D12 embryo stained for DAPI (blue), ISL1 (red), and CDX2 (green). Yellow arrows indicate the foregut and hindgut. n = 2. Scale bar, 500 μm. **i,** Representative whole-mount IF images of cultured D12 and *in vivo* E9.0 embryos stained for DAPI (blue), OCT4 (magenta), and SOX2 (yellow). Middle and lower panels show higher-magnification views of the boxed regions in the upper and middle panels, respectively. Arrows indicate primordial germ cells. n = 2 (D12) and n = 4 (E9.0). Scale bars, 500 μm (upper panels), 200 μm (middle panels), and 100 μm (lower panels). Fb, forebrain; Mb, midbrain; Hb, hindbrain; BA, branchial arches; Fg, foregut; Hg, hindgut; Ht, heart; Op, optic pit; So, somites; Tb, tail bud.
