## Extended Data Figure 1 for "Continuous *ex utero* development of mouse zygotes to early organogenesis"

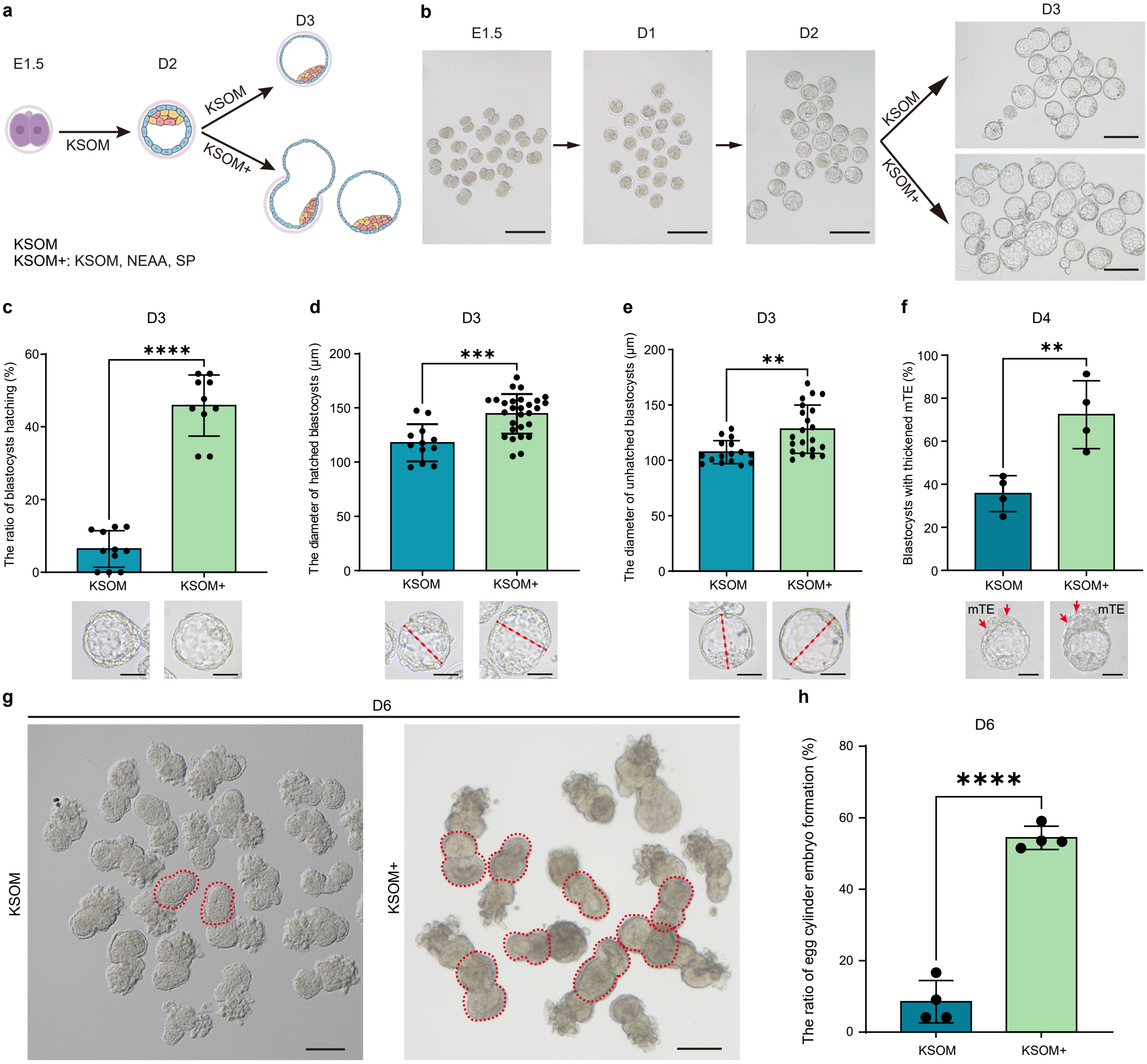
a
E1.5
D2
D3
KSOM
KSOM
KSOM+
KSOM
KSOM+: KSOM, NEAA, SP
b
E1.5
D1
D2
D3
KSOM
KSOM+
c
D3
The ratio of blastocysts hatching (%)
****
KSOM
KSOM+
d
D3
The diameter of hatched blastocysts (μm)
***
KSOM
KSOM+
e
D3
The diameter of unhatched blastocysts (μm)
**
KSOM
KSOM+
f
D4
Blastocysts with thickened mTE (%)
**
KSOM
KSOM+
mTE
mTE
g
D6
KSOM
KSOM+
h
D6
The ratio of egg cylinder embryo formation (%)
****
KSOM
KSOM+
