## Extended Data Figure 2 for "Continuous *ex utero* development of mouse zygotes to early organogenesis"

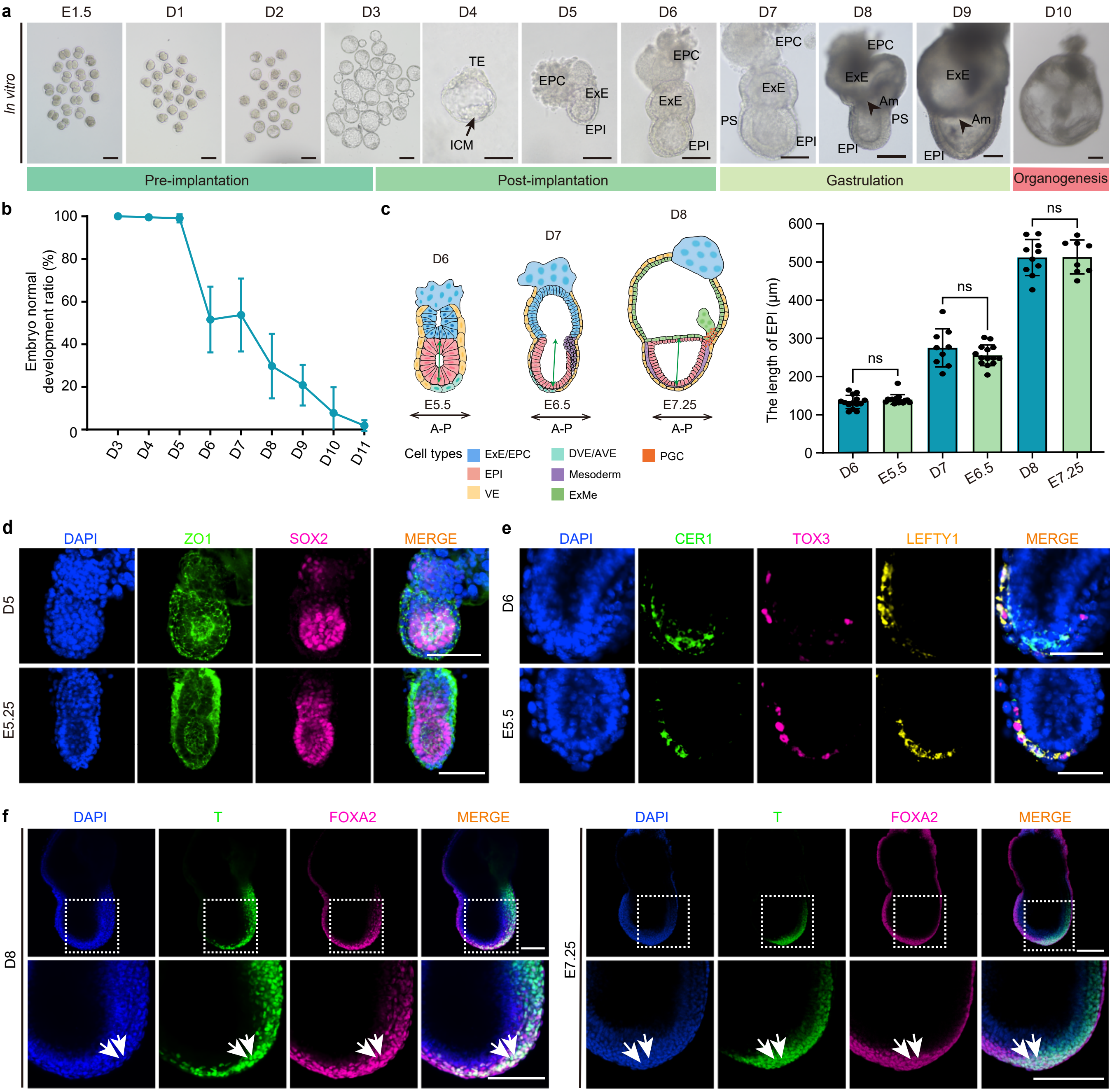
a
E1.5
D1
D2
D3
D4
D5
D6
D7
D8
D9
D10
In vitro
TE
ICM
EPC
ExE
EPI
PS
Am
Pre-implantation
Post-implantation
Gastrulation
Organogenesis
b
Embryo normal development ratio (%)
100
80
60
40
20
0
D3
D4
D5
D6
D7
D8
D9
D10
D11
c
D6
D7
D8
E5.5
E6.5
E7.25
A-P
Cell types
ExE/EPC
EPI
VE
DVE/AVE
Mesoderm
ExMe
PGC
The length of EPI (μm)
600
500
400
300
200
100
0
ns
D6
E5.5
D7
E6.5
D8
E7.25
d
DAPI
ZO1
SOX2
MERGE
D5
E5.25
e
DAPI
CER1
TOX3
LEFTY1
MERGE
D6
E5.5
f
DAPI
T
FOXA2
MERGE
D8
E7.25
