## Extended Data Figure 3 for "Continuous *ex utero* development of mouse zygotes to early organogenesis"

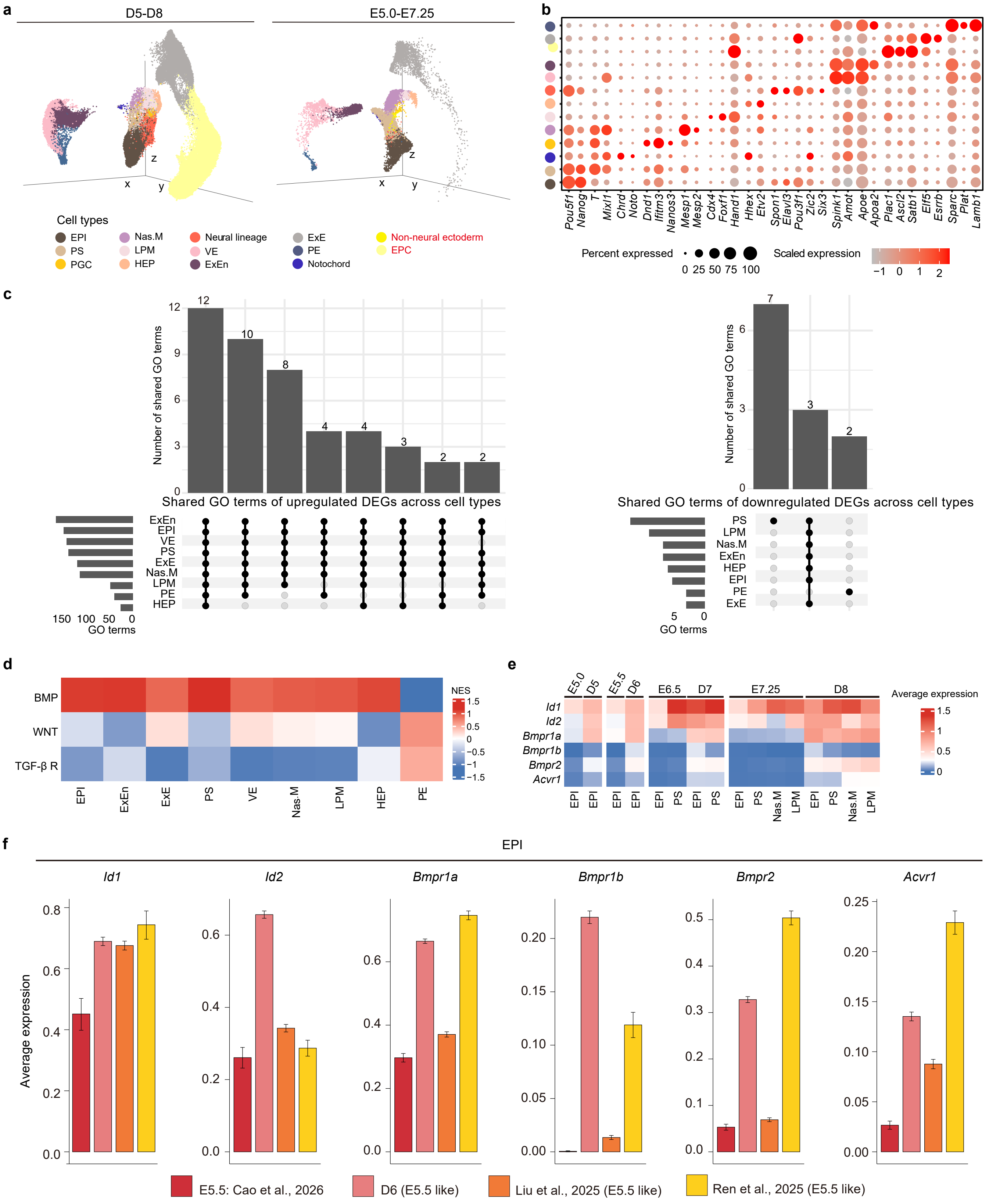

a
D5-D8
E5.0-E7.25
Cell types
EPI
PS
PGC
Nas.M
LPM
HEP
Neural lineage
VE
ExEn
ExE
PE
Notochord
Non-neural ectoderm
EPC
b
Pou5f1
Nanog
T
Mixl1
Chrd
Noto
Dnd1
Ifitm3
Nanos3
Mesp1
Mesp2
Cdx4
Foxf1
Hand1
Hhex
Etv2
Spon1
Elavl3
Pou3f1
Zic2
Six3
Spink1
Amot
Apoe
Apoa2
Plac1
Ascl2
Satb1
Elf5
Esrrb
Sparc
Plat
Lamb1
Percent expressed
0 25 50 75 100
Scaled expression
c
Number of shared GO terms
Shared GO terms of upregulated DEGs across cell types
GO terms
Shared GO terms of downregulated DEGs across cell types
d
BMP
WNT
TGF-β R
NES
e
Average expression
f
EPI
Id1
Id2
Bmpr1a
Bmpr1b
Bmpr2
Acvr1
Average expression
E5.5: Cao et al., 2026
D6 (E5.5 like)
Liu et al., 2025 (E5.5 like)
Ren et al., 2025 (E5.5 like)
