## Extended Data Figure 4 for "Continuous *ex utero* development of mouse zygotes to early organogenesis"

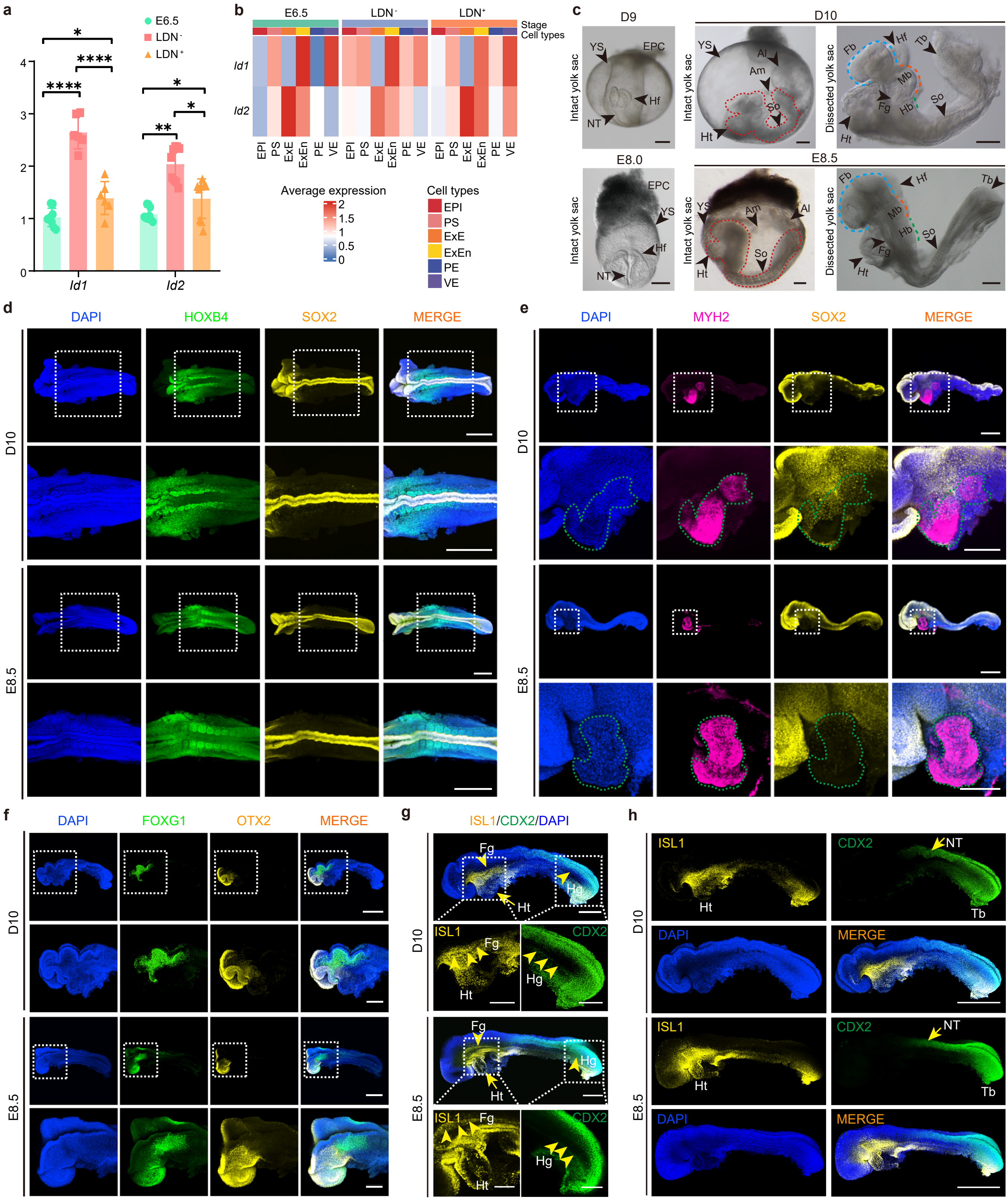
a
E6.5
LDN⁻
LDN⁺
Id1
Id2
b
E6.5
LDN⁻
LDN⁺
Stage
Cell types
Average expression
Cell types
EPI
PS
ExE
ExEn
PE
VE
c
D9
D10
E8.0
E8.5
Intact yolk sac
Dissected yolk sac
d
DAPI
HOXB4
SOX2
MERGE
D10
E8.5
e
DAPI
MYH2
SOX2
MERGE
f
DAPI
FOXG1
OTX2
MERGE
g
ISL1/CDX2/DAPI
h
ISL1
CDX2
DAPI
MERGE
