## Extended Data Figure 5 for "Continuous *ex utero* development of mouse zygotes to early organogenesis"

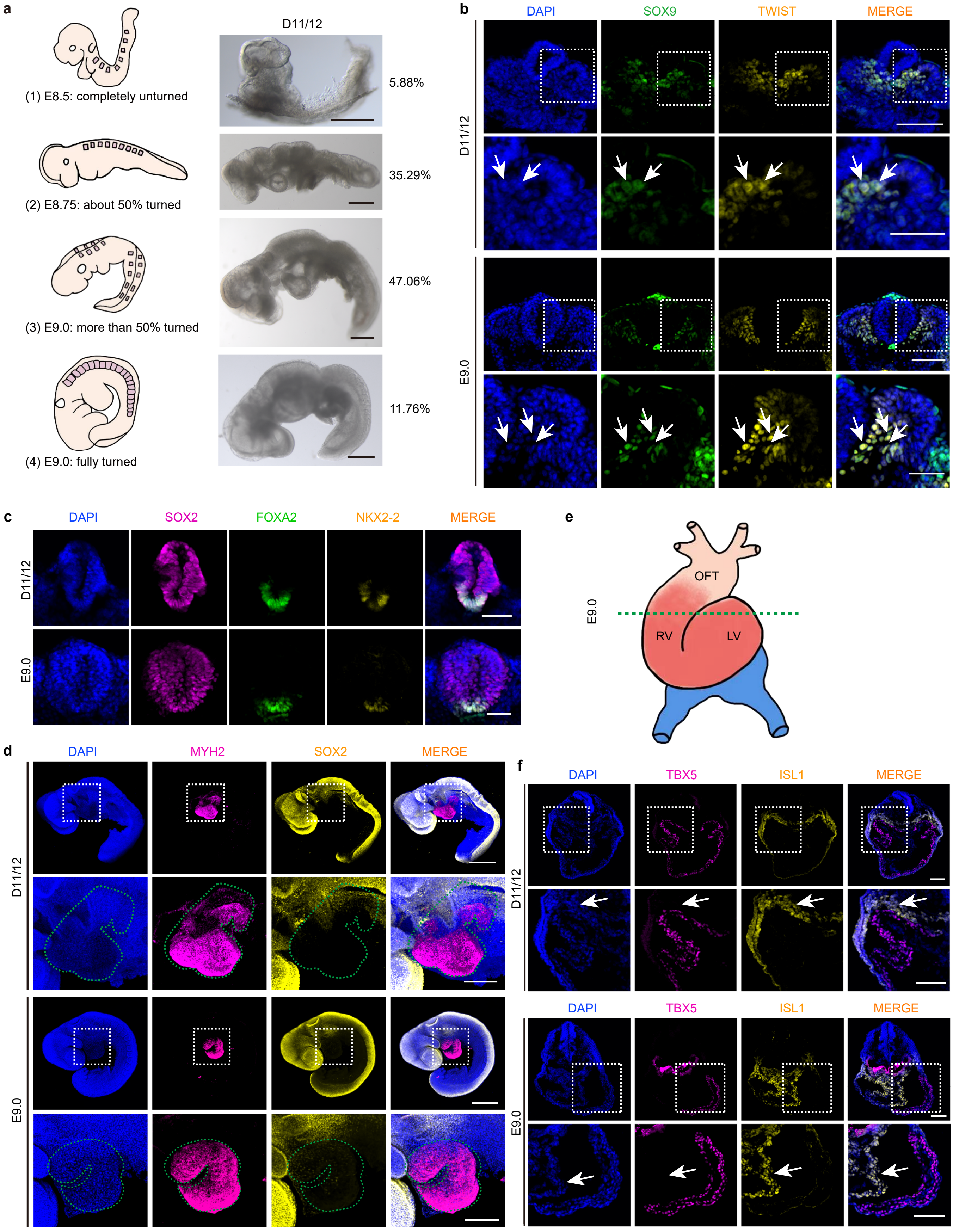
a
(1) E8.5: completely unturned
(2) E8.75: about 50% turned
(3) E9.0: more than 50% turned
(4) E9.0: fully turned
D11/12
5.88%
35.29%
47.06%
11.76%
b
DAPI
SOX9
TWIST
MERGE
D11/12
E9.0
c
DAPI
SOX2
FOXA2
NKX2-2
MERGE
D11/12
E9.0
d
DAPI
MYH2
SOX2
MERGE
D11/12
E9.0
e
E9.0
OFT
RV
LV
f
DAPI
TBX5
ISL1
MERGE
D11/12
E9.0
