## Extended Data Figure 6 for "Continuous *ex utero* development of mouse zygotes to early organogenesis"

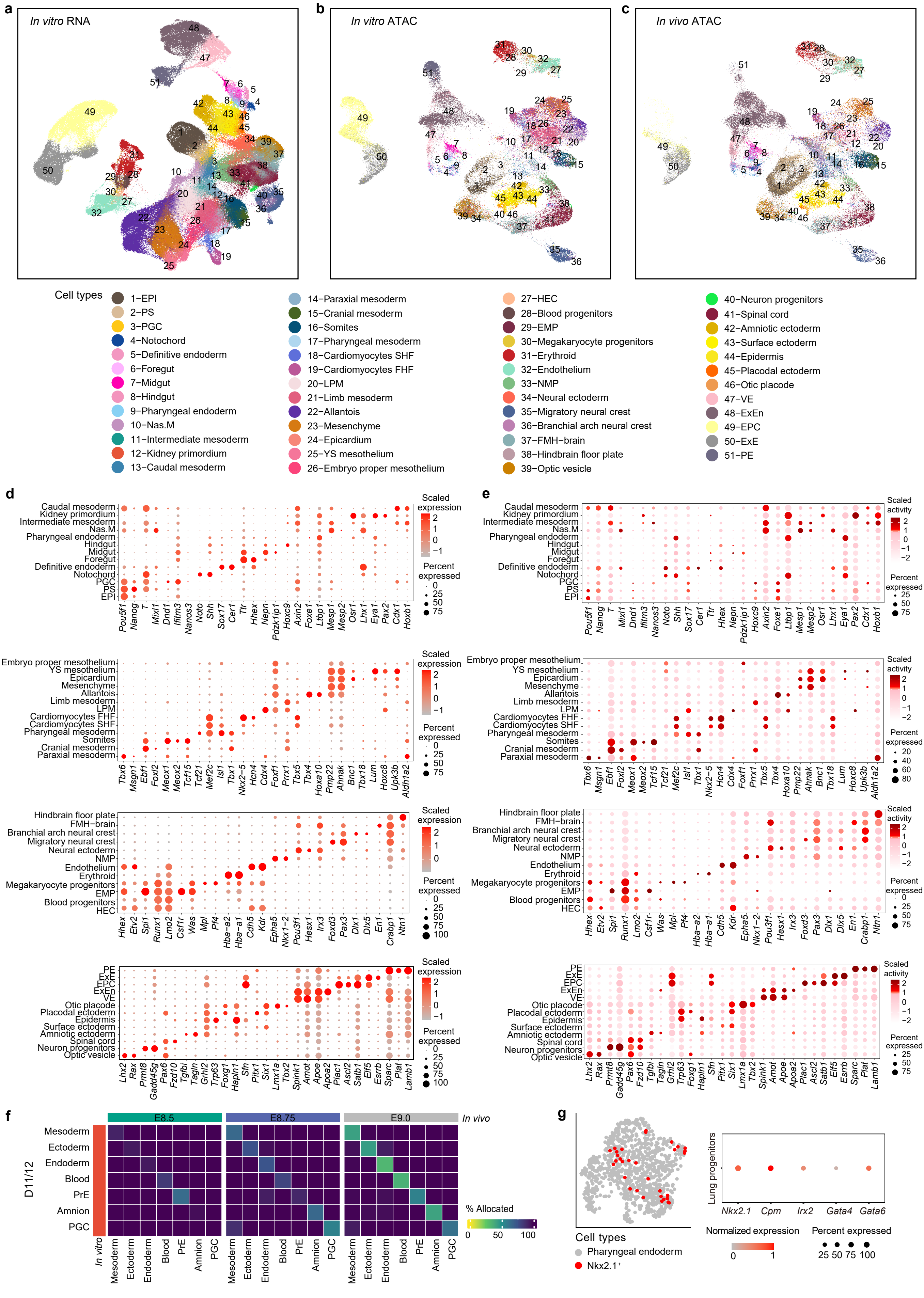

a
In vitro RNA
b
In vitro ATAC
c
In vivo ATAC
Cell types
1–EPI
2–PS
3–PGC
4–Notochord
5–Definitive endoderm
6–Foregut
7–Midgut
8–Hindgut
9–Pharyngeal endoderm
10–Nas.M
11–Intermediate mesoderm
12–Kidney primordium
13–Caudal mesoderm
14–Paraxial mesoderm
15–Cranial mesoderm
16–Somites
17–Pharyngeal mesoderm
18–Cardiomyocytes SHF
19–Cardiomyocytes FHF
20–LPM
21–Limb mesoderm
22–Allantois
23–Mesenchyme
24–Epicardium
25–YS mesothelium
26–Embryo proper mesothelium
27–HEC
28–Blood progenitors
29–EMP
30–Megakaryocyte progenitors
31–Erythroid
32–Endothelium
33–NMP
34–Neural ectoderm
35–Migratory neural crest
36–Branchial arch neural crest
37–FMH–brain
38–Hindbrain floor plate
39–Optic vesicle
40–Neuron progenitors
41–Spinal cord
42–Amniotic ectoderm
43–Surface ectoderm
44–Epidermis
45–Placodal ectoderm
46–Otic placode
47–VE
48–ExEn
49–EPC
50–ExE
51–PE
d
e
Scaled expression
Scaled activity
Percent expressed
Pou5f1 Nanog T Mixl1 Dnd1 Ifitm3 Nanos3 Noto Shh Sox17 Cer1 Ttr Hhex Nepn Pdzk1ip1 Hoxc9 Axin2 Foxe1 Ltbp1 Mesp1 Mesp2 Osr1 Lhx1 Eya1 Pax2 Cdx1 Hoxb1
Tbx6 Msgn1 Ebf1 Foxl2 Meox1 Meox2 Tcf15 Tcf21 Mef2c Isl1 Tbx1 Nkx2–5 Hcn4 Cdx4 Foxf1 Prrx1 Tbx5 Tbx4 Hoxa10 Pmp22 Ahnak Bnc1 Tbx18 Lum Hoxc8 Upk3b Aldh1a2
Hhex Etv2 Spi1 Runx1 Lmo2 Csf1r Was Mpl Pf4 Hba–a2 Hba–a1 Cdh5 Kdr Epha5 Nkx1–2 Pou3f1 Hesx1 Irx3 Foxd3 Pax3 Dlx1 Dlx5 En1 Crabp1 Ntn1
Lhx2 Rax Prmt8 Gadd45g Pax6 Fzd10 Tgfbi Tagln Grhl2 Trp63 Foxg1 Hapln1 Sfn Pitx1 Six1 Lmx1a Tbx2 Spink1 Amot Apoe Apoa2 Plac1 Ascl2 Satb1 Elf5 Esrrb Sparc Plat Lamb1
f
E8.5
E8.75
E9.0
In vivo
D11/12
In vitro
Mesoderm
Ectoderm
Endoderm
Blood
PrE
Amnion
PGC
% Allocated
g
Cell types
Pharyngeal endoderm
Nkx2.1⁺
Lung progenitors
Nkx2.1 Cpm Irx2 Gata4 Gata6
Normalized expression
Percent expressed
