## Extended Data Figure 7 for "Continuous *ex utero* development of mouse zygotes to early organogenesis"

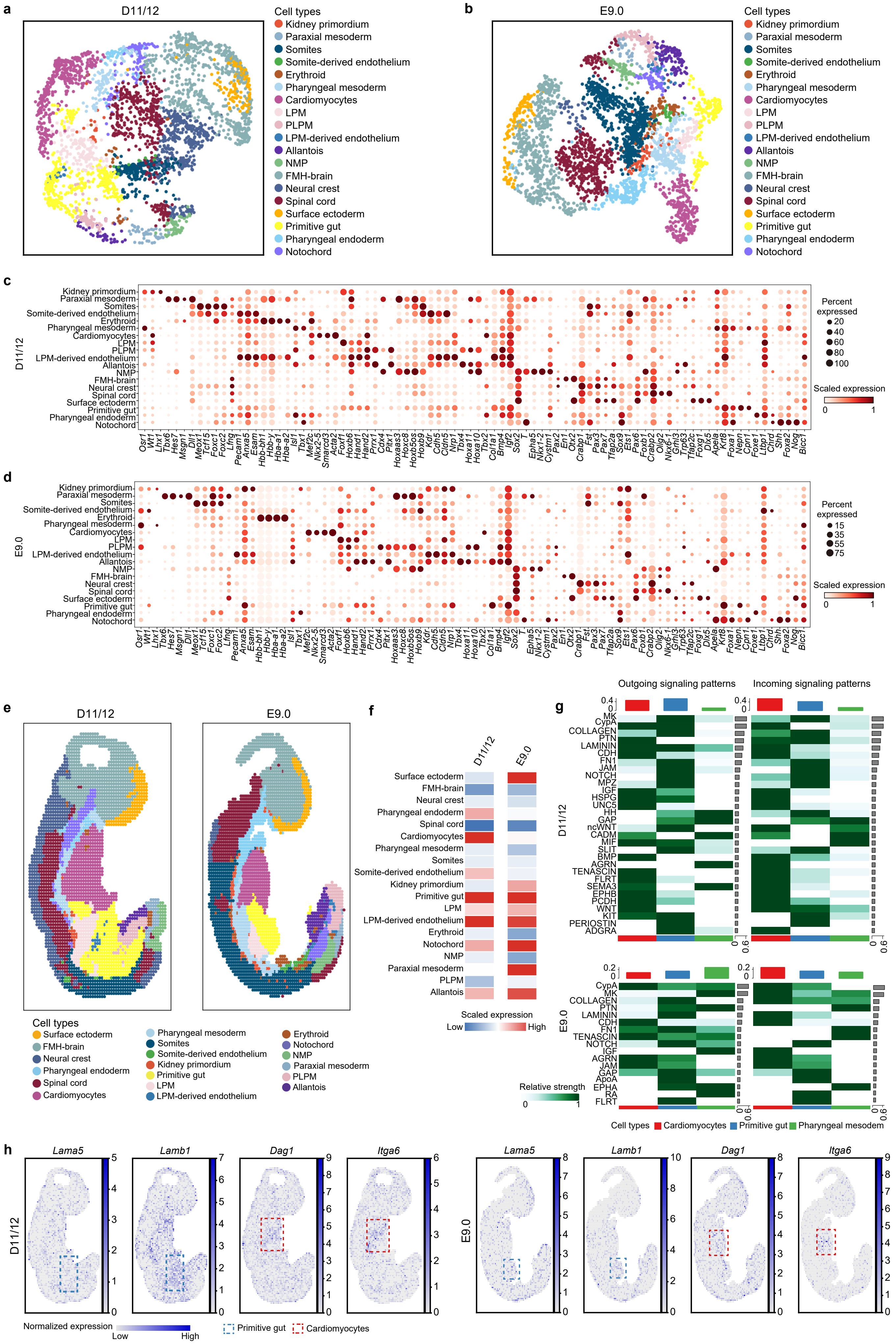
a
D11/12
b
E9.0
Cell types
Kidney primordium
Paraxial mesoderm
Somites
Somite-derived endothelium
Erythroid
Pharyngeal mesoderm
Cardiomyocytes
LPM
PLPM
LPM-derived endothelium
Allantois
NMP
FMH-brain
Neural crest
Spinal cord
Surface ectoderm
Primitive gut
Pharyngeal endoderm
Notochord
c
d
Percent expressed
Scaled expression
e
f
Scaled expression
Low
High
g
Outgoing signaling patterns
Incoming signaling patterns
Relative strength
Cell types Cardiomyocytes Primitive gut Pharyngeal mesodem
h
Lama5
Lamb1
Dag1
Itga6
Normalized expression
Low
High
Primitive gut
Cardiomyocytes
