## Extended Data Figure 8 for "Continuous *ex utero* development of mouse zygotes to early organogenesis"

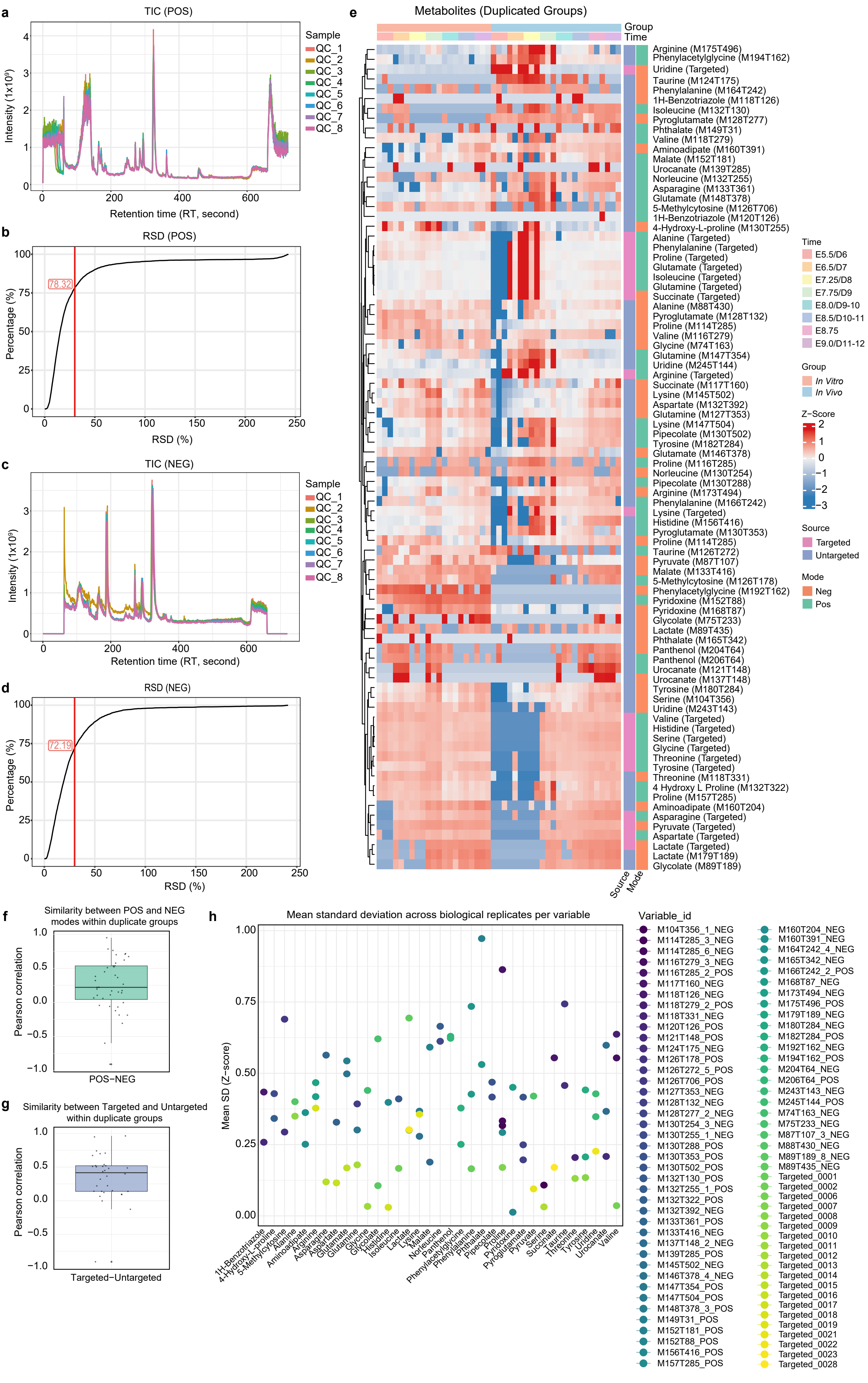
a
TIC (POS)
Intensity (1x10⁹)
Retention time (RT, second)
Sample
QC_1
QC_2
QC_3
QC_4
QC_5
QC_6
QC_7
QC_8
b
RSD (POS)
78.32
Percentage (%)
RSD (%)
c
TIC (NEG)
Intensity (1x10⁹)
Retention time (RT, second)
Sample
QC_1
QC_2
QC_3
QC_4
QC_5
QC_6
QC_7
QC_8
d
RSD (NEG)
72.19
Percentage (%)
RSD (%)
e
Metabolites (Duplicated Groups)
Group
Time
Arginine (M175T496)
Phenylacetylglycine (M194T162)
Uridine (Targeted)
Taurine (M124T175)
Phenylalanine (M164T242)
1H-Benzotriazole (M118T126)
Isoleucine (M132T130)
Pyroglutamate (M128T277)
Phthalate (M149T31)
Valine (M118T279)
Aminoadipate (M160T391)
Malate (M152T181)
Urocanate (M139T285)
Norleucine (M132T255)
Asparagine (M133T361)
Glutamate (M148T378)
5-Methylcytosine (M126T706)
1H-Benzotriazole (M120T126)
4-Hydroxy-L-proline (M130T255)
Alanine (Targeted)
Phenylalanine (Targeted)
Proline (Targeted)
Glutamate (Targeted)
Isoleucine (Targeted)
Glutamine (Targeted)
Succinate (Targeted)
Alanine (M88T430)
Pyroglutamate (M128T132)
Proline (M114T285)
Valine (M116T279)
Glycine (M74T163)
Glutamine (M147T354)
Uridine (M245T144)
Arginine (Targeted)
Succinate (M117T160)
Lysine (M145T502)
Aspartate (M132T392)
Glutamine (M127T353)
Lysine (M147T504)
Pipecolate (M130T502)
Tyrosine (M182T284)
Glutamate (M146T378)
Proline (M116T285)
Norleucine (M130T254)
Pipecolate (M130T288)
Arginine (M173T494)
Phenylalanine (M166T242)
Lysine (Targeted)
Histidine (M156T416)
Pyroglutamate (M130T353)
Proline (M114T285)
Taurine (M126T272)
Pyruvate (M87T107)
Malate (M133T416)
5-Methylcytosine (M126T178)
Phenylacetylglycine (M192T162)
Pyridoxine (M152T88)
Pyridoxine (M168T87)
Glycolate (M75T233)
Lactate (M89T435)
Phthalate (M165T342)
Panthenol (M204T64)
Panthenol (M206T64)
Urocanate (M121T148)
Urocanate (M137T148)
Tyrosine (M180T284)
Serine (M104T356)
Uridine (M243T143)
Valine (Targeted)
Histidine (Targeted)
Serine (Targeted)
Glycine (Targeted)
Threonine (Targeted)
Tyrosine (Targeted)
Threonine (M118T331)
4 Hydroxy L Proline (M132T322)
Proline (M157T285)
Aminoadipate (M160T204)
Asparagine (Targeted)
Pyruvate (Targeted)
Aspartate (Targeted)
Lactate (Targeted)
Lactate (M179T189)
Glycolate (M89T189)
Source
Mode
Time
E5.5/D6
E6.5/D7
E7.25/D8
E7.75/D9
E8.0/D9-10
E8.5/D10-11
E8.75
E9.0/D11-12
Group
In Vitro
In Vivo
Z-Score
Source
Targeted
Untargeted
Mode
Neg
Pos
f
Similarity between POS and NEG modes within duplicate groups
Pearson correlation
POS-NEG
g
Similarity between Targeted and Untargeted within duplicate groups
Pearson correlation
Targeted-Untargeted
h
Mean standard deviation across biological replicates per variable
Mean SD (Z-score)
1H-Benzotriazole
4-Hydroxy-L-proline
5-Methylcytosine
Alanine
Aminoadipate
Arginine
Asparagine
Aspartate
Glutamate
Glutamine
Glycine
Glycolate
Histidine
Isoleucine
Lactate
Lysine
Malate
Norleucine
Panthenol
Phenylacetylglycine
Phenylalanine
Phthalate
Pipecolate
Proline
Pyridoxine
Pyroglutamate
Pyruvate
Serine
Succinate
Taurine
Threonine
Tyrosine
Uridine
Urocanate
Valine
Variable_id
M104T356_1_NEG
M114T285_3_NEG
M114T285_6_NEG
M116T279_3_NEG
M116T285_2_POS
M117T160_NEG
M118T126_NEG
M118T279_2_POS
M118T331_NEG
M120T126_POS
M121T148_POS
M124T175_NEG
M126T178_POS
M126T272_5_POS
M126T706_POS
M127T353_NEG
M128T132_NEG
M128T277_2_NEG
M130T254_3_NEG
M130T255_1_NEG
M130T288_POS
M130T353_POS
M130T502_POS
M132T130_POS
M132T255_1_POS
M132T322_POS
M132T392_NEG
M133T361_POS
M133T416_NEG
M137T148_2_NEG
M139T285_POS
M145T502_NEG
M146T378_4_NEG
M147T354_POS
M147T504_POS
M148T378_3_POS
M149T31_POS
M152T181_POS
M152T88_POS
M156T416_POS
M157T285_POS
M160T204_NEG
M160T391_NEG
M164T242_4_NEG
M165T342_NEG
M166T242_2_POS
M168T87_NEG
M173T494_NEG
M175T496_POS
M179T189_NEG
M180T284_NEG
M182T284_POS
M192T162_NEG
M194T162_POS
M204T64_NEG
M206T64_POS
M243T143_NEG
M245T144_POS
M74T163_NEG
M75T233_NEG
M87T107_3_NEG
M88T430_NEG
M89T189_8_NEG
M89T435_NEG
Targeted_0001
Targeted_0002
Targeted_0006
Targeted_0007
Targeted_0008
Targeted_0009
Targeted_0010
Targeted_0011
Targeted_0012
Targeted_0013
Targeted_0014
Targeted_0015
Targeted_0016
Targeted_0017
Targeted_0018
Targeted_0019
Targeted_0021
Targeted_0022
Targeted_0023
Targeted_0028
