## Extended Data Figure 9 for "Continuous *ex utero* development of mouse zygotes to early organogenesis"

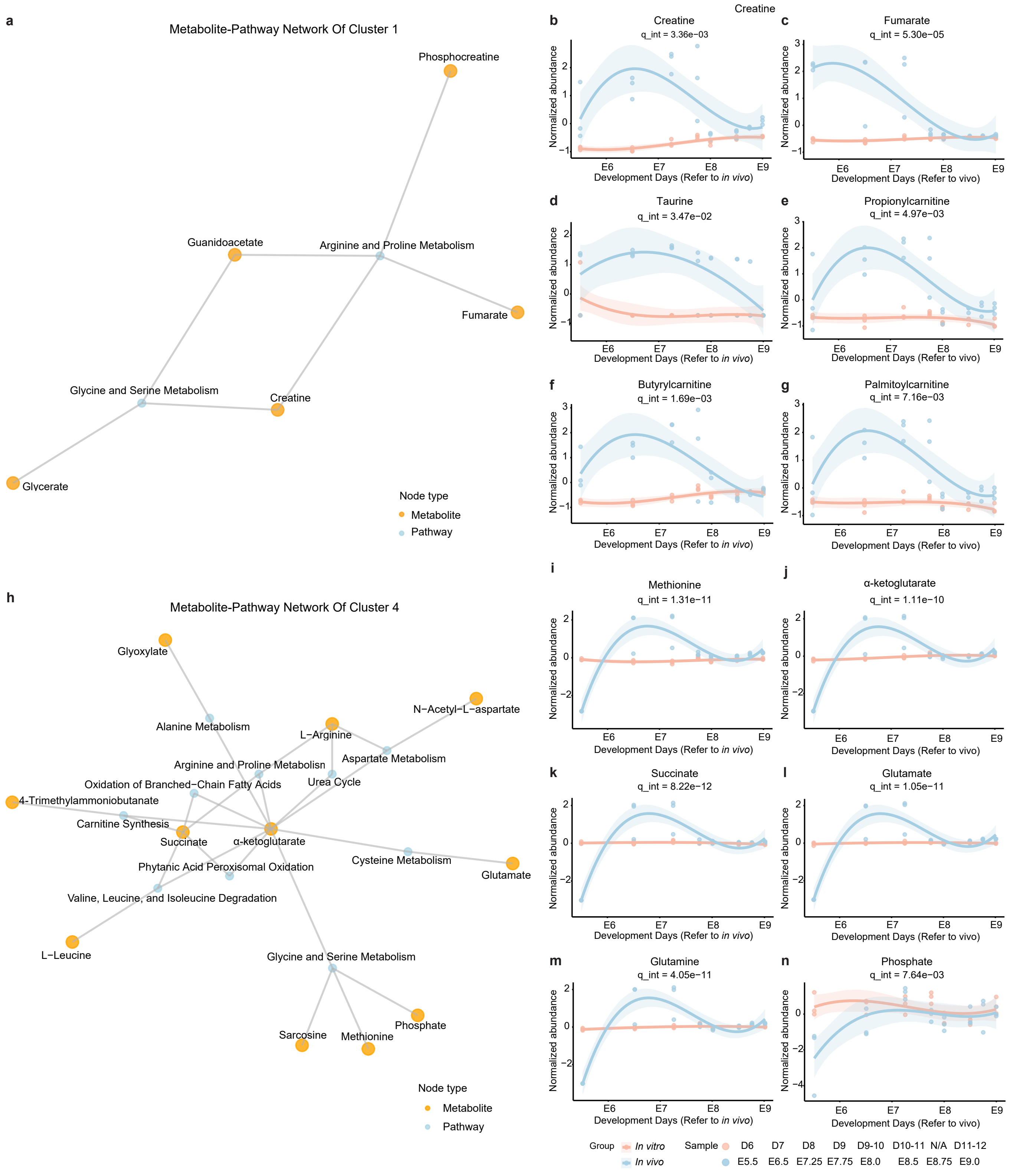

Creatine
a
Metabolite-Pathway Network Of Cluster 1
Phosphocreatine
Guanidoacetate
Arginine and Proline Metabolism
Fumarate
Glycine and Serine Metabolism
Creatine
Glycerate
Node type
Metabolite
Pathway
b
Creatine
q_int = 3.36e−03
c
Fumarate
q_int = 5.30e−05
d
Taurine
q_int = 3.47e−02
e
Propionylcarnitine
q_int = 4.97e−03
f
Butyrylcarnitine
q_int = 1.69e−03
g
Palmitoylcarnitine
q_int = 7.16e−03
Normalized abundance
Development Days (Refer to in vivo)
Development Days (Refer to vivo)
E6
E7
E8
E9
h
Metabolite-Pathway Network Of Cluster 4
Glyoxylate
Alanine Metabolism
L−Arginine
N−Acetyl−L−aspartate
Arginine and Proline Metabolism
Aspartate Metabolism
Oxidation of Branched−Chain Fatty Acids
Urea Cycle
4-Trimethylammoniobutanate
Carnitine Synthesis
Succinate
α-ketoglutarate
Cysteine Metabolism
Glutamate
Phytanic Acid Peroxisomal Oxidation
Valine, Leucine, and Isoleucine Degradation
L−Leucine
Glycine and Serine Metabolism
Sarcosine
Methionine
Phosphate
Node type
Metabolite
Pathway
i
Methionine
q_int = 1.31e−11
j
α-ketoglutarate
q_int = 1.11e−10
k
Succinate
q_int = 8.22e−12
l
Glutamate
q_int = 1.05e−11
m
Glutamine
q_int = 4.05e−11
n
Phosphate
q_int = 7.64e−03
Group
In vitro
In vivo
Sample
D6 D7 D8 D9 D9-10 D10-11 N/A D11-12
E5.5 E6.5 E7.25 E7.75 E8.0 E8.5 E8.75 E9.0
