## Extended Data Figure 10 for "Continuous *ex utero* development of mouse zygotes to early organogenesis"

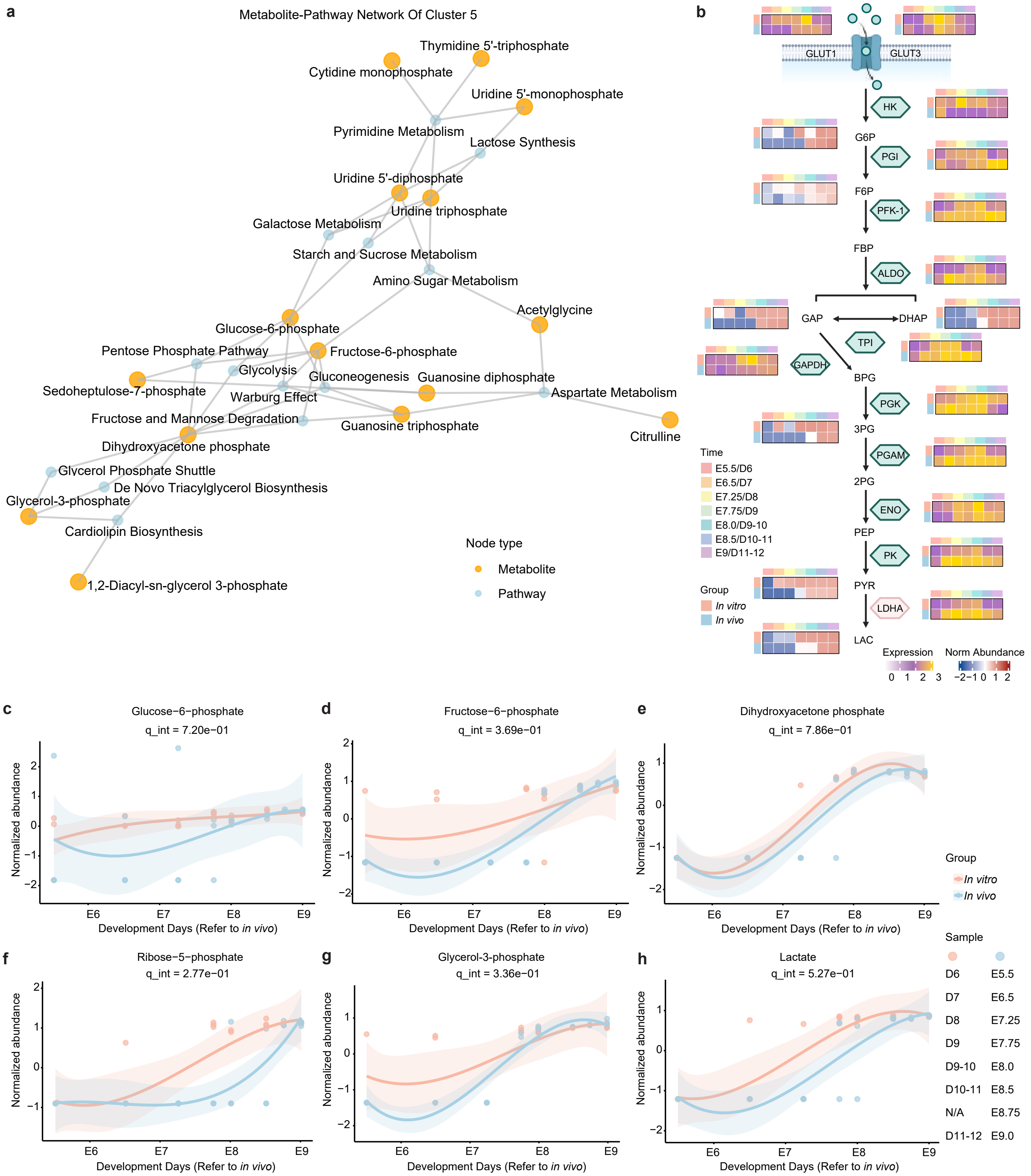
a
Metabolite-Pathway Network Of Cluster 5
Thymidine 5'-triphosphate
Cytidine monophosphate
Uridine 5'-monophosphate
Pyrimidine Metabolism
Lactose Synthesis
Uridine 5'-diphosphate
Uridine triphosphate
Galactose Metabolism
Starch and Sucrose Metabolism
Amino Sugar Metabolism
Acetylglycine
Glucose-6-phosphate
Pentose Phosphate Pathway
Fructose-6-phosphate
Glycolysis
Gluconeogenesis
Guanosine diphosphate
Sedoheptulose-7-phosphate
Warburg Effect
Aspartate Metabolism
Fructose and Mannose Degradation
Guanosine triphosphate
Citrulline
Dihydroxyacetone phosphate
Glycerol Phosphate Shuttle
De Novo Triacylglycerol Biosynthesis
Glycerol-3-phosphate
Cardiolipin Biosynthesis
1,2-Diacyl-sn-glycerol 3-phosphate
Node type
Metabolite
Pathway
b
GLUT1
GLUT3
HK
G6P
PGI
F6P
PFK-1
FBP
ALDO
GAP
DHAP
TPI
GAPDH
BPG
PGK
3PG
PGAM
2PG
ENO
PEP
PK
PYR
LDHA
LAC
Time
E5.5/D6
E6.5/D7
E7.25/D8
E7.75/D9
E8.0/D9-10
E8.5/D10-11
E9/D11-12
Group
In vitro
In vivo
Expression
0 1 2 3
Norm Abundance
−2 −1 0 1 2
c
Glucose−6−phosphate
q_int = 7.20e−01
d
Fructose−6−phosphate
q_int = 3.69e−01
e
Dihydroxyacetone phosphate
q_int = 7.86e−01
f
Ribose−5−phosphate
q_int = 2.77e−01
g
Glycerol-3-phosphate
q_int = 3.36e−01
h
Lactate
q_int = 5.27e−01
Normalized abundance
E6 E7 E8 E9
Development Days (Refer to in vivo)
Group
In vitro
In vivo
Sample
D6 E5.5
D7 E6.5
D8 E7.25
D9 E7.75
D9-10 E8.0
D10-11 E8.5
N/A E8.75
D11-12 E9.0
