## Extended Data Figure 11 for "Continuous *ex utero* development of mouse zygotes to early organogenesis"

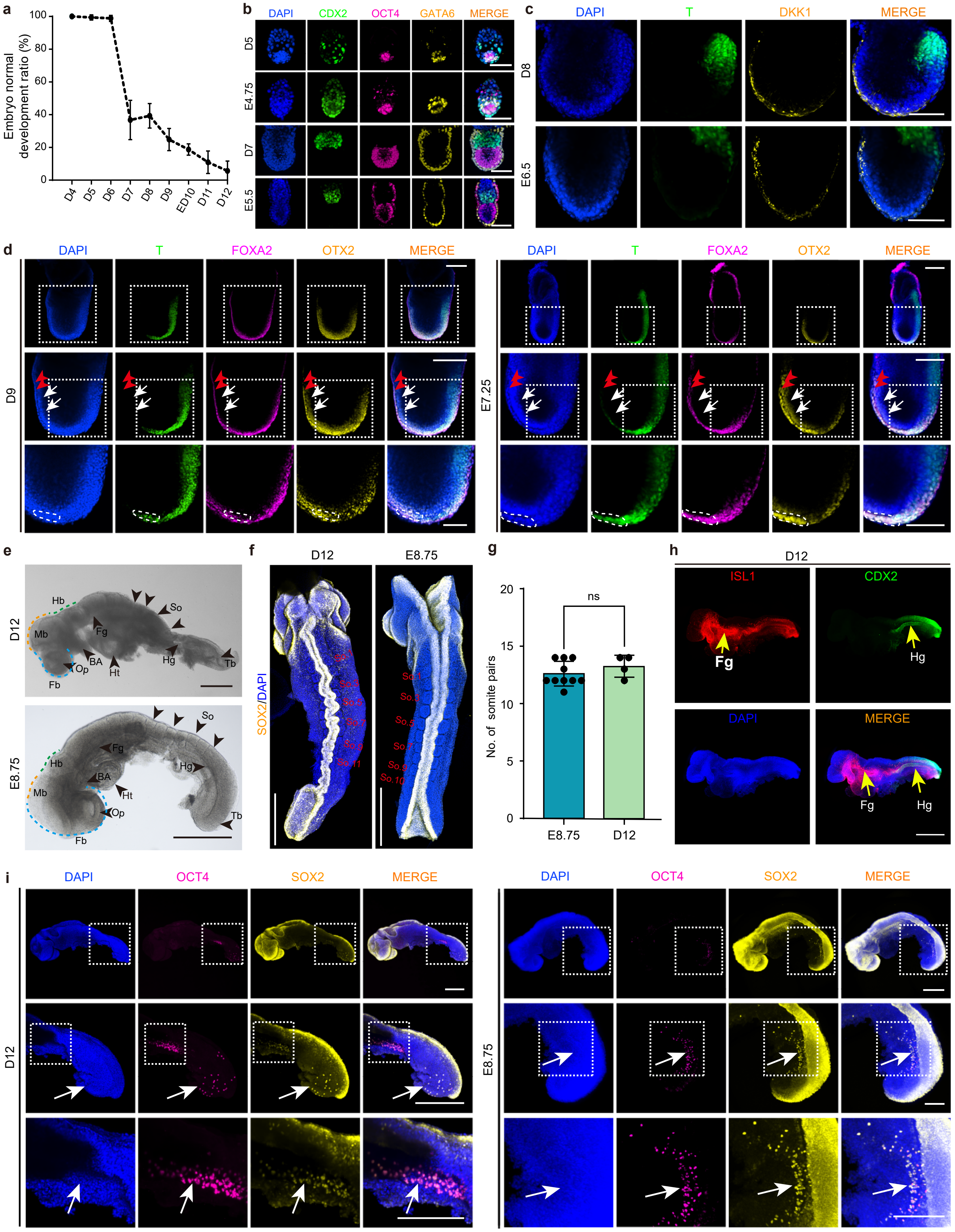
a
Embryo normal development ratio (%)
D4
D5
D6
D7
D8
D9
ED10
D11
D12
b
DAPI
CDX2
OCT4
GATA6
MERGE
D5
E4.75
D7
E5.5
c
DAPI
T
DKK1
MERGE
D8
E6.5
d
DAPI
T
FOXA2
OTX2
MERGE
D9
E7.25
e
D12
E8.75
Hb
Mb
Fb
Op
BA
Ht
Fg
Hg
So
Tb
f
D12
E8.75
SOX2/DAPI
g
ns
No. of somite pairs
E8.75
D12
h
D12
ISL1
CDX2
DAPI
MERGE
Fg
Hg
i
DAPI
OCT4
SOX2
MERGE
D12
E8.75
